# Improved CUT&RUN chromatin profiling and analysis tools

**DOI:** 10.1101/569129

**Authors:** Michael P. Meers, Terri Bryson, Steven Henikoff

## Abstract

We previously described a novel alternative to Chromatin Immunoprecipitation, Cleavage Under Targets & Release Using Nuclease (CUT&RUN), in which unfixed permeabilized cells are incubated with antibody, followed by binding of a Protein A-Micrococcal Nuclease (pA/MNase) fusion protein (1). Upon activation of tethered MNase, the bound complex is excised and released into the supernatant for DNA extraction and sequencing. Here we introduce four enhancements to CUT&RUN: 1) a hybrid Protein A-Protein G-MNase construct that expands antibody compatibility and simplifies purification; 2) a modified digestion protocol that inhibits premature release of the nuclease-bound complex; 3) a calibration strategy based on carry-over of *E. coli* DNA introduced with the fusion protein; and 4) a novel peak-calling strategy customized for the low-background profiles obtained using CUT&RUN. These new features, coupled with the previously described low-cost, high efficiency, high reproducibility and high-throughput capability of CUT&RUN make it the method of choice for routine epigenomic profiling.

## Introduction

Profiling the chromatin landscape for specific components is one of the most widely used methods in biology, and over the past decade, chromatin immunoprecipitation (ChIP) followed by sequencing (ChIP-seq) has become practically synonymous with chromatin profiling (2, 3). However, the most widely used ChIP-seq protocols have limitations and are subject to artifacts (4-7), of which only some have been addressed by methodological improvements (8-12). An inherent limitation to ChIP is that solubilization of chromatin, whether by sonication or enzymatic digestion, results in sampling from the entire solubilized genome, and this requires very deep sequencing so that the sites of targeted protein binding can be resolved above background (2). To overcome this limitation, we introduced Cleavage Under Targets and Release Using Nuclease (CUT&RUN) (1), which is based on the chromatin immunocleavage (ChIC) targeted nuclease strategy (13): Successive incubation of unfixed cells or nuclei with an antibody and a Protein A-Micrococcal Nuclease (pA/MNase) fusion protein is followed by activation of MNase with calcium. In CUT&RUN, cells or nuclei remain intact throughout the procedure and only the targeted sites of binding are released into solution. Our CUT&RUN method dramatically reduced non-specific backgrounds, such that ∼10-fold lower sequencing depth was required to obtain similar peak-calling performance (1). In addition, CUT&RUN provides near base-pair resolution, and our most recently published benchtop protocol is capable of profiling ∼100 human cells for an abundant histone modification and ∼1000 cells for a transcription factor (14). The simplicity of CUT&RUN has also resulted in a fully automated robotic version (AutoCUT&RUN) in which the high reproducibility and low cost makes it ideally suited for high-throughput epigenomic profiling of clinical samples (15). Other advances based on our original CUT&RUN publication include CUT&RUN.Salt for fractionation of chromatin based on solubility (16) and CUT&RUN.ChIP for profiling specific protein components within complexes released by CUT&RUN digestion (17). CUT&RUN has also been adopted by others (18-32), and since publication of our *eLife* paper we have distributed materials to >500 laboratories world-wide, with user questions and answers fielded interactively on our open-access Protocols.io site (33).

Broad implementation of CUT&RUN requires reagent and bioinformatics standardization, and the rapid adoption of CUT&RUN by the larger community of researchers motivates the enhancements described here. First, the method requires a fusion protein that is not at this writing commercially available, and the published pA/MNase purification protocol is cumbersome, which effectively restricts dissemination of the method. Therefore we have produced an improved construct with a 6-His-Tag that can be easily purified using a commercial kit, and by using a Protein A-Protein G hybrid, the fusion protein binds avidly to mouse antibodies, which bind only weakly to Protein A. Second, the original protocols are sensitive to digestion time, in that under-digestion results in low yield and over-digestion can result in pre-mature release of pA/MNase-bound complexes that can digest accessible DNA sites. To address this limitation, we have modified the protocol such that premature release is reduced, allowing digestion to near-completion for high yields with less background. Third, the current CUT&RUN protocol recommends a spike-in of heterologous DNA at the release step to compare samples in a series. Here we demonstrate that adding a spike-in is unnecessary, because the carry-over of *E. coli* DNA from purification of pA/MNase or pAG/MNase is sufficient to calibrate samples in a series. Finally, popular peak-calling algorithms designed for ChIP-seq data are crippled by the reduced background noise in CUT&RUN experiments, so we introduce a novel peak-caller that takes advantage of sparse background to define peaks and set thresholds, and we show that it provides better performance on CUT&RUN data than two widely used peak-calling algorithms on both narrow peaks and broad domains.

## Results and Discussion

### An improved CUT&RUN vector

The pA/MNase fusion protein produced by the pK19-pA-MN plasmid (13) requires purification from lysates of *Escherichia coli* overexpressing cells using an immunoglobulin G (IgG) column, and elution with low pH followed by neutralization has resulted in variations between batches. To improve the purification protocol, we added a 6-His tag (34) into the pK19-pA-MN fusion protein (Figure 1A and Figure 1 – figure supplement 1A). This allowed for simple and gentle purification on a nickel resin column (Figure 1 – figure supplement 1B). In addition, we found that a commercial 6-His-cobalt resin kit also yielded pure highly active enzyme from a 20 ml culture, enough for ∼10,000 reactions. Even when used in excess, there is no increase in release of background fragments (Figure 1 – figure supplement 2), which indicates that the washes are effective in removing unbound fusion protein.

**Figure 1.**
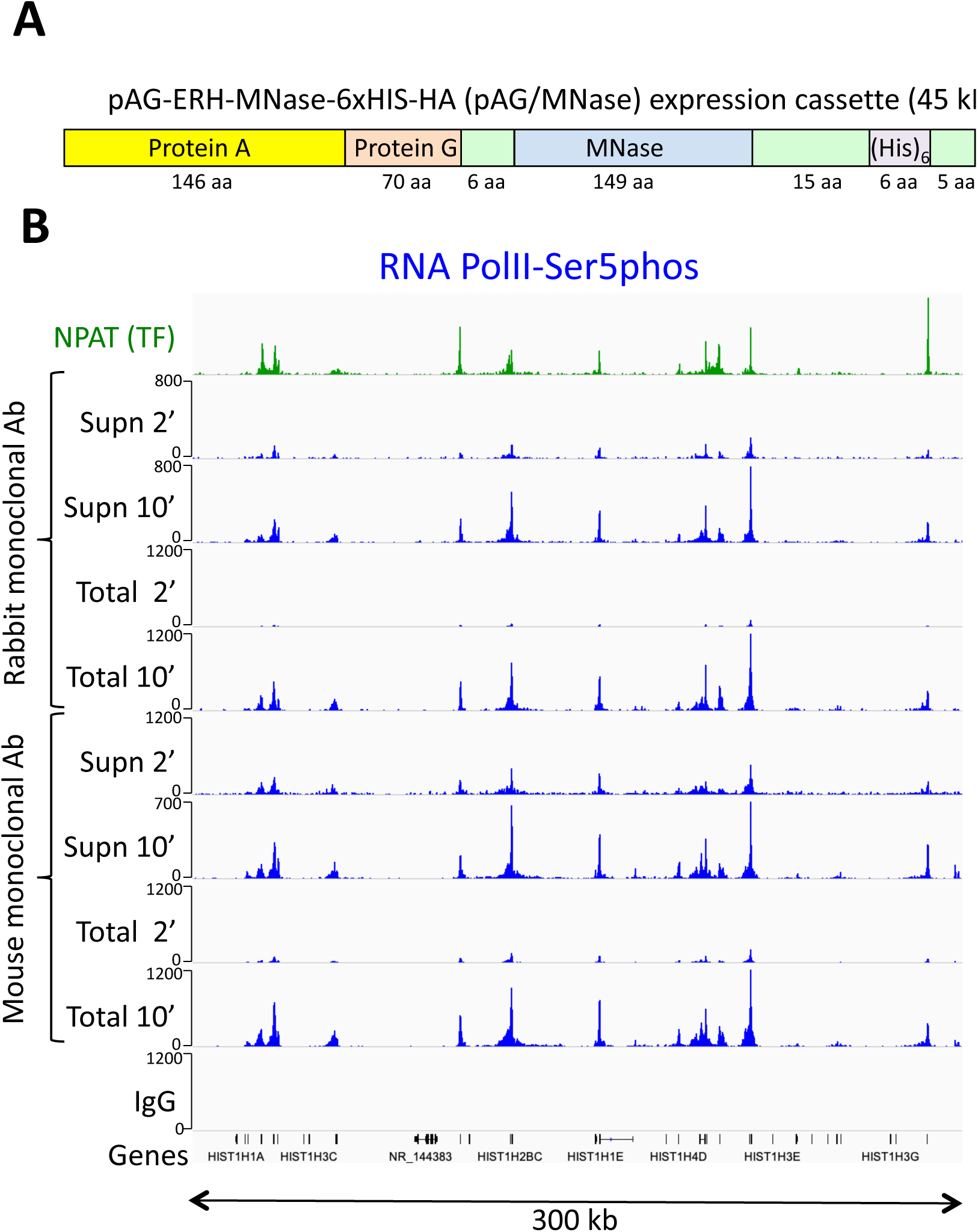
An improved fusion protein for CUT&RUN. **A**) Schematic diagram (not to scale) showing improvements to the pA-MNase fusion protein, which include addition of the C2 Protein G IgG binding domain, a 6-histidine tag for purification and a hemagglutinin tag (HA) for immunoprecipitation. **B**) The Protein A/G hybrid fusion results in high-efficiency CUT&RUN for both rabbit and mouse primary antibodies. CUT&RUN for both rabbit and mouse RNAPII-Ser5phosphate using pAG/MNase were extracted from either the supernatant or the total cellular extract. Tracks are shown for the histone gene cluster at Chr6:26,000,000-26,300,000, where NPAT is a transcription factor that co-activates histone genes. Tracks for 2’ and 10’ time points are displayed at the same scale for each antibody and for both supernatant (supn) or total DNA extraction protocols.

In principle an epitope tagged pAG/MNase could be used for chromatin pull-down from a CUT&RUN supernatant in sequential strategies like CUT&RUN.ChIP (17). However, in practice use of the 6-His tag is complicated by the requirement for a chelating agent to release the protein from the nickel resin. Therefore, we also added an HA (hemagglutinin) tag, which could be used to affinity-purify the complex of a directly bound chromatin particle with a primary antibody and the fusion protein.

Protein A binds well to rabbit, goat, donkey and guinea pig IgG antibodies, but poorly to mouse IgG1, and so for most mouse antibodies, Protein G is generally used (35). To further improve the versatility of the MNase fusion protein, we encoded a single Protein G domain adjacent to the Protein A domain in the pK19-pA-MN plasmid (36). In addition, we mutated three residues in the Protein G coding sequence to further increase binding for rabbit antibodies (37). This resulted in a fusion protein that binds strongly to most commercial antibodies without requiring a secondary antibody. We found that for ordinary CUT&RUN applications pAG/MNase behaves very similarly to pA/MNase, but is more easily purified and is more versatile, for example allowing us to perform CUT&RUN without requiring a secondary antibody for mouse primary monoclonal antibodies (Figure 1B).

### Preventing premature release during CUT&RUN digestion

When fragments are released by cleavage in the presence of Ca^++^ ions, the associated pA/MNase complex can digest accessible DNA (1). Although performing digestion at 0°C minimizes this artifact, eliminating premature release during digestion would allow for more complete release of target-specific fragments. Based on the observation that nucleosome core particles aggregate in high-divalent-cation and low-salt conditions (38), we wondered whether these conditions would prevent premature release of chromatin particles *in situ*. Therefore, we performed digestions in 10 mM CaCl_2_ and 3.5 mM HEPES pH 7.5. Under these high-calcium/low-salt conditions, chromatin is digested with no detectable release of fragments into the supernatant (Figure 2). Reactions are halted by transferring the tube to a magnet, removing the liquid, and adding elution buffer containing 150 mM NaCl, 20 mM EGTA and 25 µg/ml RNAse A, which releases the small DNA fragments into the supernatant. These conditions are compatible with direct end-polishing and ligation used for AutoCUT&RUN (15). Furthermore, retention of the cleaved fragments within the nucleus under high-divalent cation/low-salt conditions could facilitate single-cell application of CUT&RUN.

**Figure 2.**
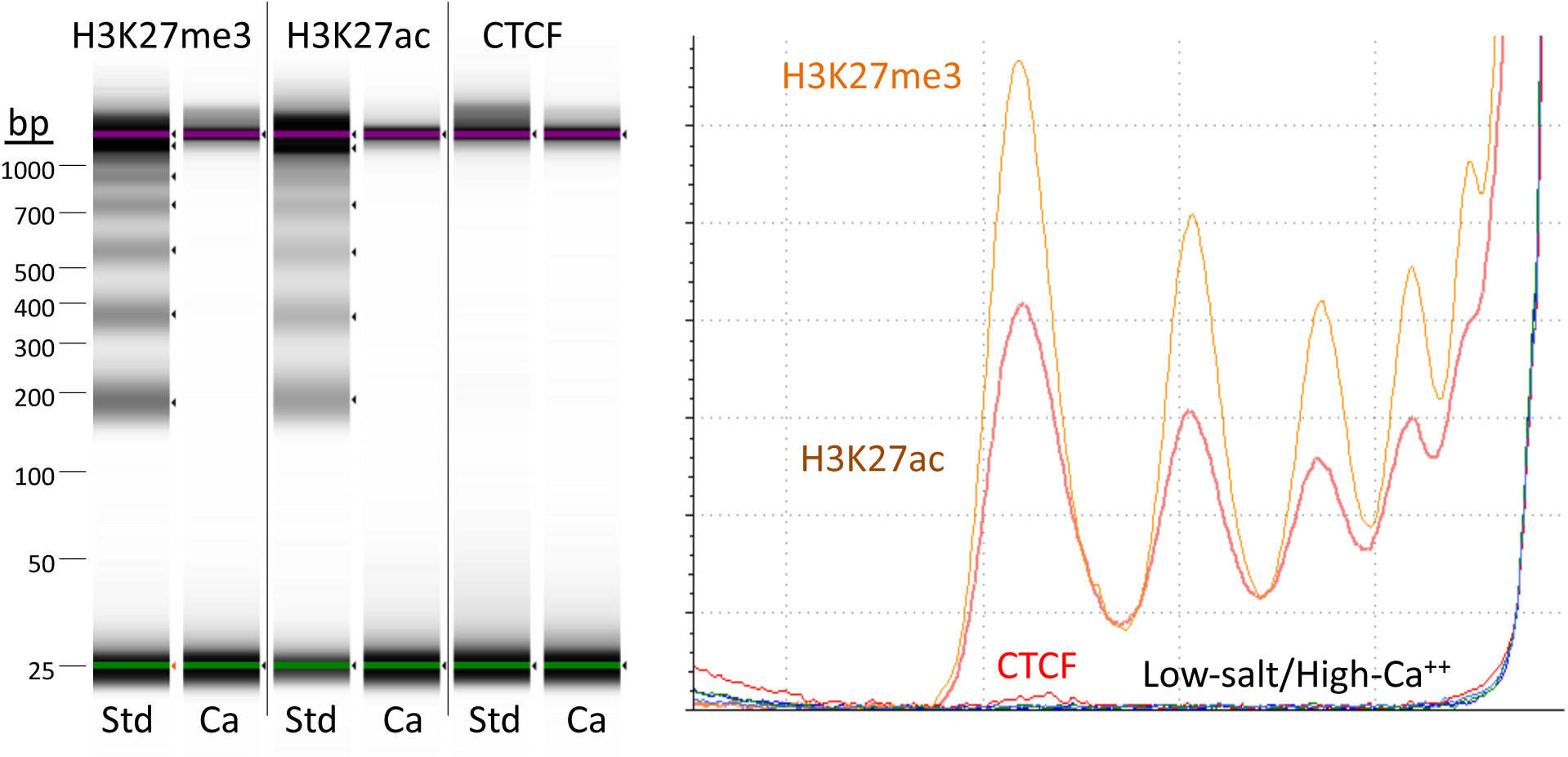
Targeted fragments are not released during digestion using high-calcium/low-salt conditions. CUT&RUN was performed using either the high-Ca^++^/low-salt (Ca^++^) or the standard (Std) method with antibodies to three different epitopes. DNA was extracted from supernatants, where no elution was carried out for the Ca^++^ samples. Although high yields of nucleosomal ladder DNA eluted from the supernatants using the standard method, no DNA was detectable in the supernatant using the high-Ca^++^/low salt method when the elution step was omitted. Left, Tapestation images from indicated lanes; Right, Densitometry of the same lanes.

The high-calcium/low-salt protocol provided similar results using either pA/MNase and pAG/MNase (Figure 3). We also obtained similar results with either protocol for digestion time points over a ∼30-fold range and for both supernatant and total DNA extraction (Figure 4 – figure supplement 1). For antibodies to H3K27ac, libraries produced using the high-calcium/low-salt protocol showed improved consistency relative to the standard protocol when digested over an extended time-course (Figure 4), presumably because preventing release of particles during digestion avoids their premature release where they would artifactually digest accessible DNA. The close correlations between high-calcium/low-salt H3K27ac datasets for time points over a ∼100-fold range occur with corresponding increases in the yield of fragments released into the supernatant during subsequent elution (Figure 4 – figure supplement 2). This indicates that longer digestion times result in higher yields, with high signal-to-noise throughout the digestion series (Figure 4). Thus, this modification of CUT&RUN can reduce the risk of overdigestion for abundant epitopes such as H3K27ac, where premature release of pA-MNase-bound chromatin particles can increase background.

**Figure 3.**
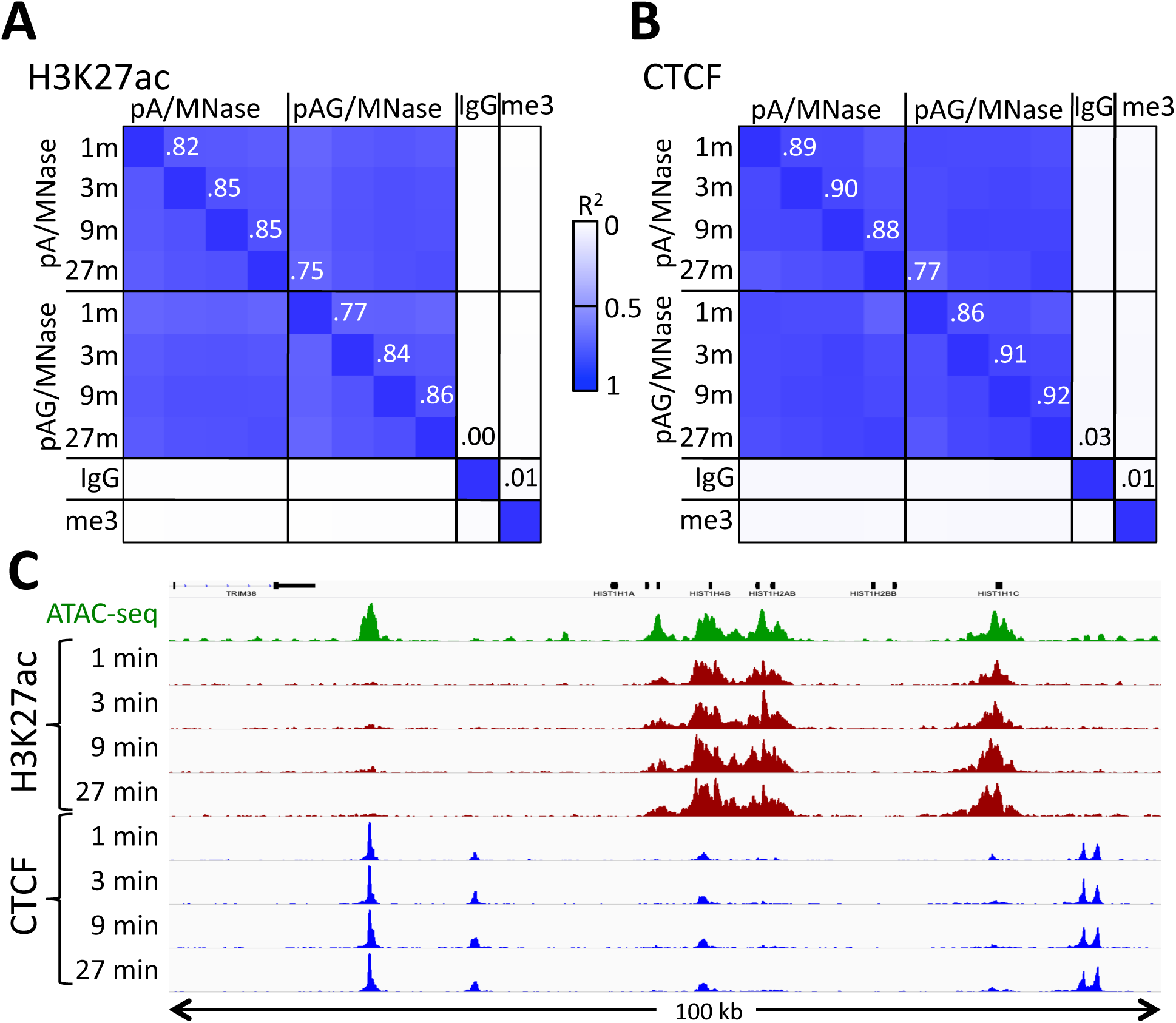
Similar performance using pA/MNase and pAG/MNase. **A**) CUT&RUN was performed with an antibody to H3K27ac (Millipore MABE647) and to CTCF (Millipore 07-729) with digestion over a 1 to 27 min range as indicated using pA/G-MNase with the high-Ca^++^/low-salt protocol. Correlation matrix comparing peak overlap for time points and fusion constructs. The datasets were pooled and MACS2 was used with default parameters to call peaks, excluding those in repeat-masked intervals and those where peaks overlapped with the top 1% of IgG occupancies, for a total of 52,425 peaks. Peak positions were scored for each dataset and correlations (R^2^ values shown along the diagonal and displayed with Java TreeView v.1.16r2, contrast = 1.25) were calculated between peak vectors. IgG and H3K27me3 (me3) negative controls were similarly scored. **B**) Same as A, except the antibody was to CTCF. A set of 9,403 sites with a CTCF motif within a hypersensitive site was used (Skene & Henikoff, 2017). High correlations between all time points demonstrate the uniformity of digestion over a 27-fold range. **C)** Representative tracks from datasets used for panels A and B showing a 100-kb region that includes a histone locus cluster (chr6:25,972,600-26,072,600).

**Figure 4.**
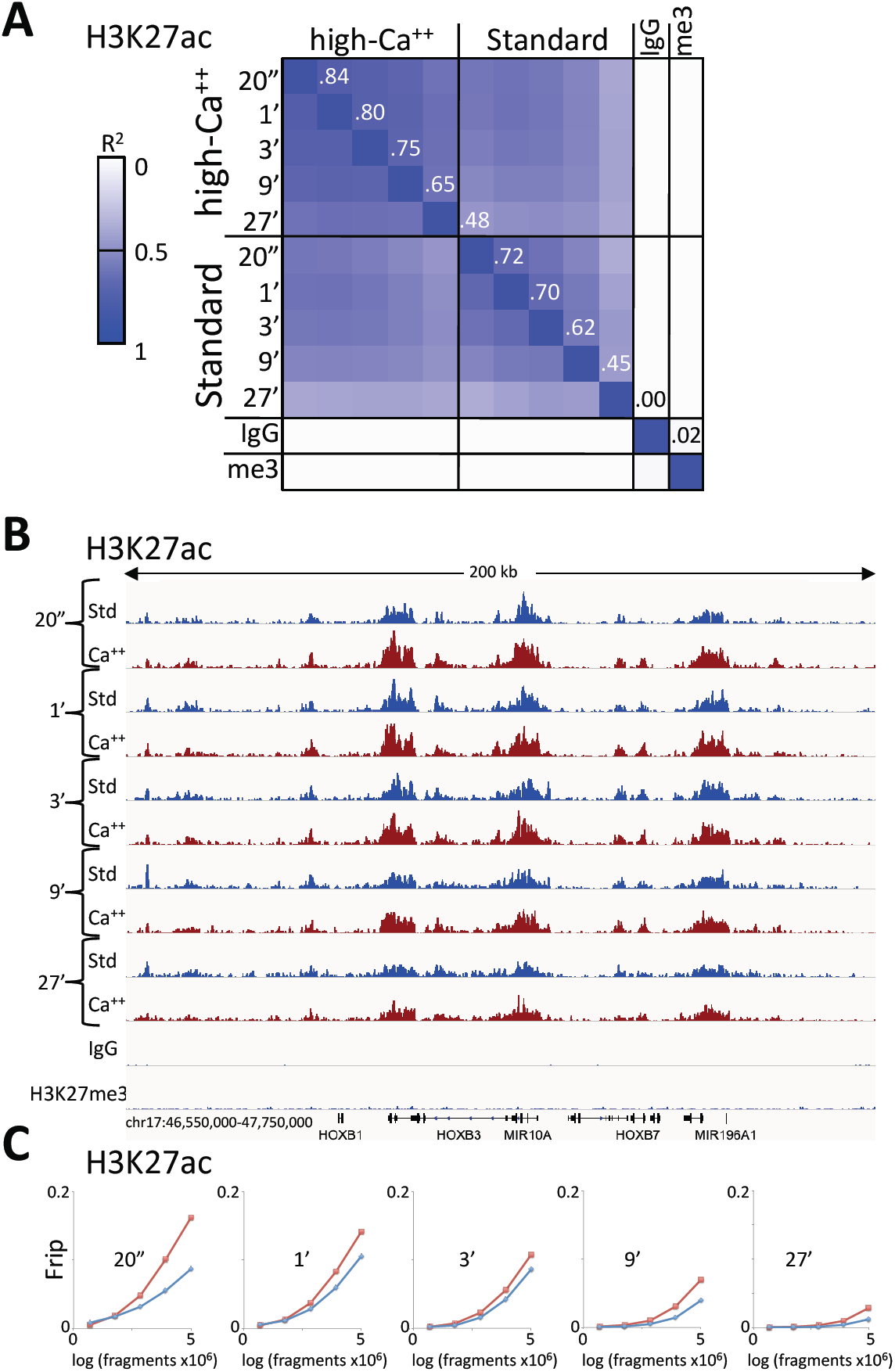
Consistent peak definition with high-Ca++/low salt digestion. **A**) H3K27ac CUT&RUN time-course experiments were performed with an Abcam 4729 rabbit polyclonal antibody, following either the standard protocol or the low-salt/high-calcium (High-Ca^++^) protocol. A sample of 5 million fragments from each of the 10 H3K27ac datasets were pooled and MACS2 called 36,529 peaks. Peak positions were scored for each dataset and correlations (R^2^ values shown along the diagonal) were calculated between peak vectors. IgG and H3K27me3 (me3) negative controls were similarly scored. Higher correlations between the High-Ca^++^ than the Standard time points indicates improved uniformity of digestion over the ∼100-fold range of digestion times. **B**) Tracks from a representative 200 kb region around the HoxB locus. **C**) Fraction of reads in peaks (Frip) plots for each time point after down-sampling (5 million, 2.5 million, 1.25 million, 625,000 and 312,500).

We previously showed that CUT&RUN can be performed on insoluble protein complexes by extracting total DNA (1) or by performing salt fractionation of the bead-bound cells and extracting DNA from the residual pellet (16). In either case, large DNA fragments were depleted using SPRI (AMPure XP) beads before library preparation. RNA polymerase II (RNAPII) from animal cells is insoluble when engaged (39, 40), and requires harsh treatments for quantitative profiling using ChIP (41). To determine whether CUT&RUN can be used for insoluble chromatin complexes, we profiled Serine-5-phosphate on the C-terminal domain (CTD) of the Rpb1 subunit of RNAPII using both extraction of supernatant and of total DNA. This CTD phosphorylation is enriched in the initiating form of RNAPII, and we observed similar genic profiles for supernatant and total DNA (Figure 1B).

### Calibration using *E. coli* carry-over DNA

Comparing samples in a series typically requires calibration for experimental quality and sequencing read depth. It is common to use background levels to calibrate ChIP-seq samples in a series and to define and compare peaks for peak-calling (2). However, the extremely low backgrounds of CUT&RUN led us to a calibration strategy based on spike-in of heterologous DNA, which has been generally recommended for all situations in which samples in a series are to be compared (42, 43). In our current spike-in protocol, the heterologous DNA, which is typically DNA purified from an MNase digest of yeast *Saccharomyces cerevisiae* or *Drosophila melanogaster* chromatin, is added when stopping a reaction, and we adopted this spike-in procedure for the high-calcium/low-salt protocol described in the previous section. Interestingly, we noticed that mapping reads to both the spike-in genome and the *E. coli* genome resulted in almost perfect correlation (R^2^=0.97) between *S. cerevisiae* and *E. coli* in an experiment using pA/MNase in which the number of cells was varied over several orders of magnitude (Figure 5A). Near-perfect correlations (R^2^=0.96-0.99) between yeast spike-in and carry-over *E. coli* DNA were also seen in series using the same batch of pAG/MNase with high-calcium/low-salt digestion conditions (Figure 5B), and for both supernatant release and extraction and total DNA extraction (Figure 5C-D). These strong positive correlations are not accounted for by cross-mapping of the yeast spike-in to the *E. coli* genome, because omitting the spike-in for a low-abundance epitope resulted in very few yeast counts with high levels of *E. coli* counts (blue symbol in Figure 5C-D panels). As the source of *E. coli* DNA is carried over from purification of pA/MNase and pAG/MNase, the close correspondence provides confirmation of the accuracy of our heterologous spike-in procedure (1). Moreover, as carry-over *E. coli* DNA is introduced at an earlier step, and is cleaved to small mappable fragments that are released during digestion and elution, it provides a more desirable calibration standard than using heterologous DNA (42, 43). High correlations were also seen between *S. cerevisiae* spike-in and *E. coli* carry-over DNA for pA-MNase in batches that we have distributed (Figure 5 – table supplement 1). Therefore, data for nearly all CUT&RUN experiments performed thus far can be recalibrated *post-hoc* whether or not a spike-in calibration standard had been added.

**Figure 5.**
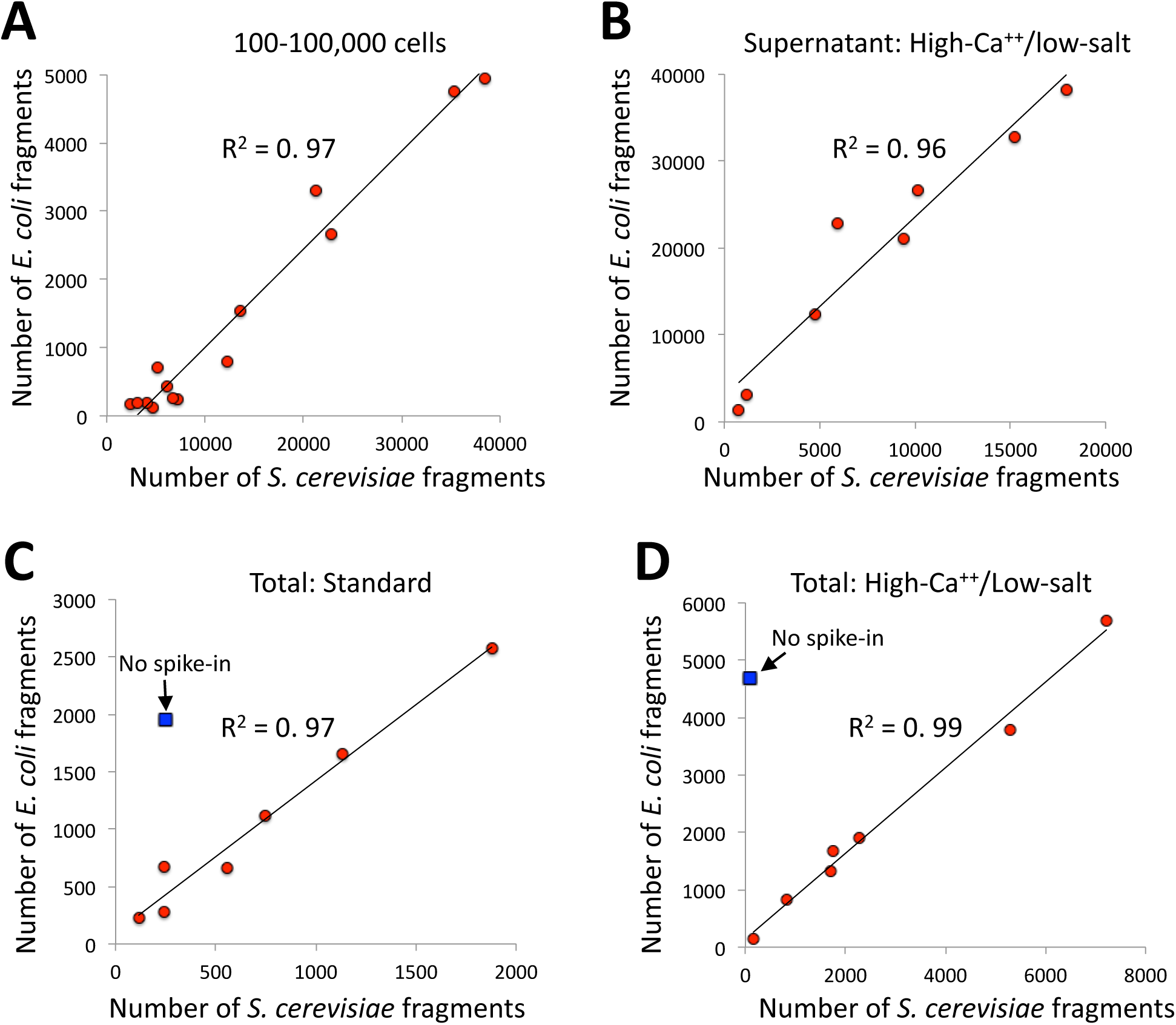
*E. coli* carry-over DNA of pA/MNase and pAG/MNase can substitute for spike-in calibration. **A**) Fragments from a CUT&RUN K562 cell experiment (GSE104550 20170426) using antibodies against H3K27me3 (100-8,000 cells) and CTCF (1,000-100,000 cells) were mapped to the repeat-masked genome of *S. cerevisae* and the full genome *E. coli*. Standard digestion was followed by supernatant release and extraction. **B**) Same as A using antibodies against multiple epitopes of varying abundances, high-calcium/low-salt digestion and supernatant release and extraction. **C**) Same as B except using standard digestion conditions and total DNA extraction. The *S.* cerevisiae spike-in DNA was left out for one sample (blue square). From top to bottom, antibodies are: NPAT Thermo PA5-66839, Myc: CST Rabbit Mab #13987, CTD: PolII CTD Abcam 8WG16, RNAPII-Ser5: Abcam 5408 (mouse), RNAPII-Ser2: CST E1Z3G, CTCF Millipore 07-729, RNAPII-Ser5: CST D9N5I (rabbit), H3K4me2: Upstate 07-030. **D**) Same as C except using high-calcium/low-salt digestion and total DNA extraction. From top to bottom, antibodies are: CTCF Millipore 07-729, NPAT Thermo PA5-66839, Myc: CST Rabbit Mab #13987, CTD: PolII CTD Abcam 8WG16, RNAPII-Ser5: Abcam 5408 (mouse), RNAPII-Ser5: CST D9N5I (rabbit), RNAPII-Ser2: CST E1Z3G, H3K4me2: Upstate 07-030.

### Peak-calling based on fragment block aggregation

Peak calling algorithms designed for the analysis of ChIP-seq data are often optimized for high recall to distinguish signal from background (44). However, the low read depths of CUT&RUN data render standard peak callers vulnerable to reduced precision (i.e. avoidance of false positives) due to the sparseness of the background, resulting in any spurious background read being called as a peak. To address this problem we developed Sparse Enrichment Analysis for CUT&RUN (SEACR), a peak-calling algorithm that enforces precision from sparse data by quantifying the global distribution of background signal and using it to set a stringent empirical threshold for peak identity. CUT&RUN data from target antibody and IgG control experiments are first parsed into “signal blocks” representing segments of continuous, non-zero read depth, and the signal in each block is calculated by summing read counts (Figure 6A). A plot of the proportion of signal blocks in Target or IgG (y-axis) is used to identify the threshold value at which the percentage of Target versus IgG blocks is maximized; then target blocks failing to meet the threshold are filtered out, leaving enriched “peaks” (Figure 6A). We also filtered out blocks that overlap an IgG block meeting the threshold as a means to eliminate spurious peaks that arise either through multiple mapping at repeated regions or by “hyper-accessibility” (4). Since SEACR is model-free and empirically data driven, it does not require arbitrary selection of parameters from a statistical model on the part of the user.

**Figure 6.**
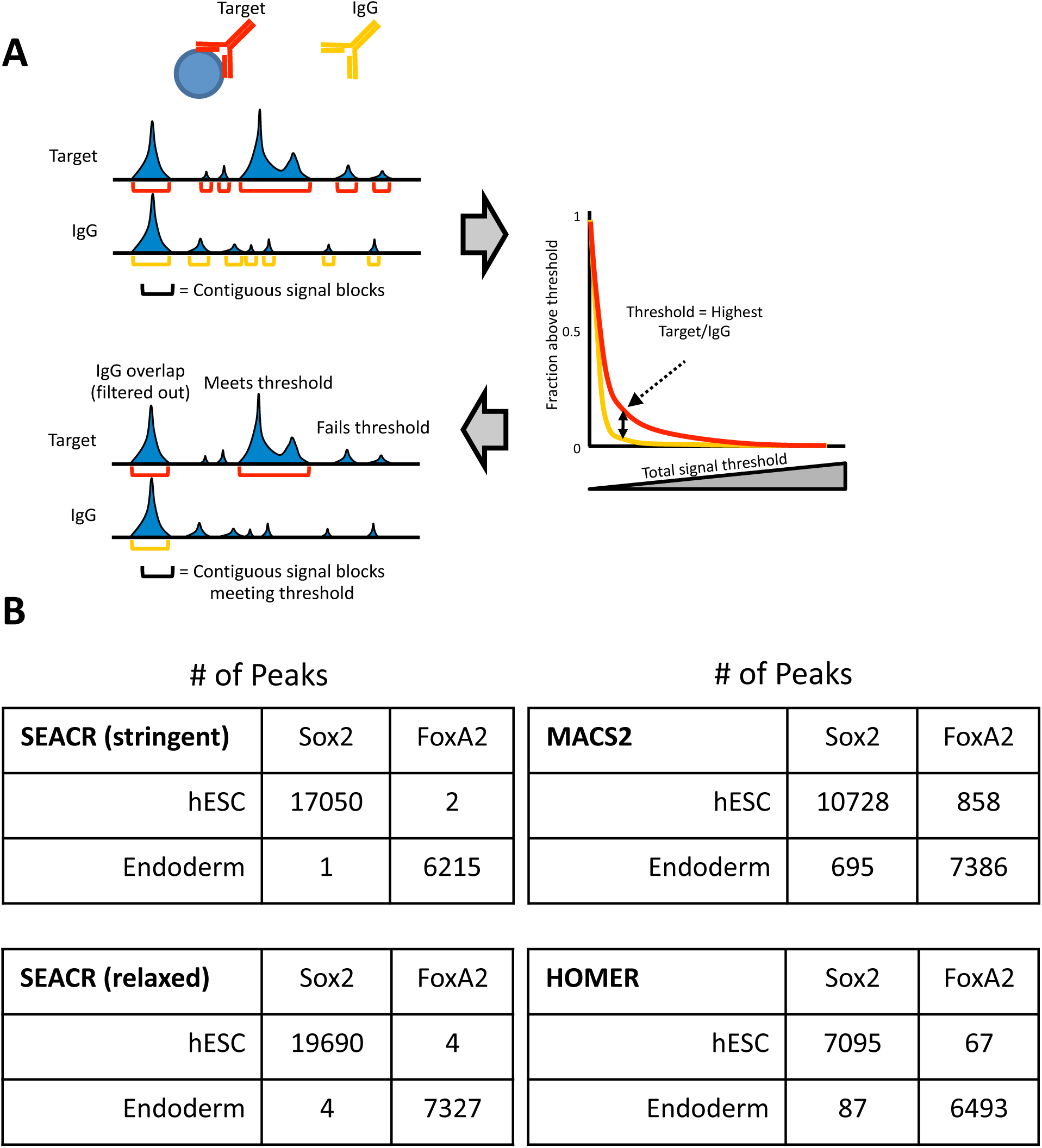
SEACR enforces peak calling specificity across a range of read depths. **A)** Schematic of SEACR methodology. Contiguous signal blocks (top) are identified and plotted by the percentage of blocks exceeding a total signal threshold (right), and an optimal threshold is empirically identified and used for filtering (bottom). **B)** Peaks were called from Sox2 or FoxA2 experiments carried out in either hESCs or Endoderm cells using SEACR in stringent or relaxed mode; MACS2; or HOMER.

To test the performance of SEACR in comparison with popular peak calling algorithms designed for ChIP-seq, we used SEACR, MACS2 and HOMER to call peaks from CUT&RUN data for two transcription factors (TFs), Sox2 and FoxA2 in human embryonic stem cells (hESCs) and Definitive Endoderm (DE) cells. Sox2 expression is restricted to hESCs and FoxA2 expression is restricted to DE cells. SEACR succeeded in calling a comparable number of peaks for Sox2 in hESCs or FoxA2 in DE cells as either MACS2 (45) or HOMER (46), while calling only 1-4 peaks for either factor when they are not expressed (Figure 6B, “stringent”). In contrast, both HOMER and MACS2 called up to ∼900 spurious peaks in these datasets; these trends held when analyzing total bases covered by peaks or percentage of reads in peaks (Figure 6—figure supplement 1A-B). We also sought to increase the recall (i.e. detection of true positives) of SEACR by including peaks that meet a maximum signal threshold (i.e. the highest read depth in the signal block) that may not have met the total signal threshold described above. We found that this approach yielded more peaks with nearly identical precision, and therefore designated this a “relaxed” mode for peak calling, in contrast with the “stringent” default (Figure 6B, Figure 4—figure supplement 1A-B, “relaxed”).

To test SEACR performance across a spectrum of read depths, we called peaks using SEACR relaxed mode (owing to its improved recall and comparable precision), MACS2 in “narrow peak” mode, and HOMER in “factor” mode, from H3K4me2 CUT&RUN data subsampled 10 times each at 11 different read depths spanning from 2 million to 40 million reads. We then called peaks and compared them with peaks called by the ENCODE consortium using MACS2 on ENCODE ChIP-seq data, assigning ENCODE peaks meeting a −log_10_ FDR threshold of greater than 10 as a stringent “truth set”. SEACR consistently minimized the fraction of called peaks that were outside the test set (false positive rate, Figure 7A-C). This indicates that the precision of SEACR is robust across a range of read depths. Precision at 30 million and 40 million reads were notable exceptions, which likely owes to the total signal threshold becoming undermined by high background. Indeed, when we implement a genome coverage threshold that converts regions of very low read coverage to 0 such that at least 50% of the reference genome contains 0 signal, SEACR shows better precision than MACS2 and HOMER above 30 million reads (Figure 7—figure supplement 1A).

**Figure 7.**
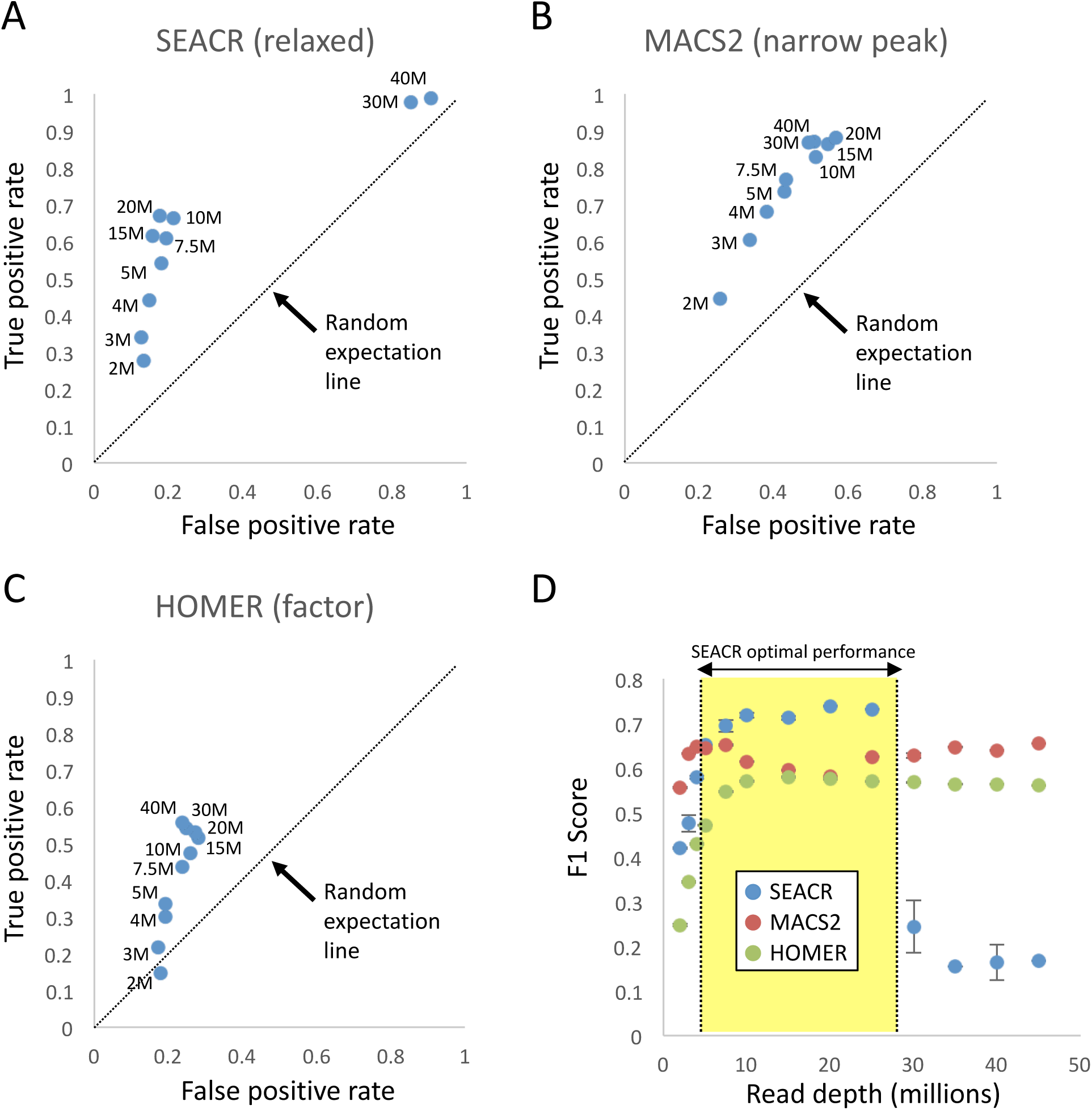
SEACR peak-calling provides an improved specificity/sensitivity trade-off. Fraction of ENCODE peaks called from subsampled H3K4me2 CUT&RUN data using SEACR relaxed mode (**A**), MACS2 narrow peak mode (**B**), or HOMER factor mode (**C**). Subsampling level is indicated next to each point. **D)** F1 score for each point on the plots in A-C; read depths at which SEACR performance is superior are indicated by yellow shaded region.

SEACR maintained higher recall (true positive rate) than HOMER across most read subsampling levels, as measured by the percentage of test set peaks overlapping CUT&RUN peaks called by the algorithm; however, in several cases recall was less than that of MACS2 at the same read depth level. To analyze the general optimality of combined recall and precision for each peak caller, we calculated the F1 score for each peak caller at each read subsampling level, such that larger F1 scores corresponded with higher performance in a combination of the two metrics. SEACR exhibited superior performance at all subsampling levels between 5M and 25M reads (Figure 7D). To account for the fact that peak callers such as MACS2 have parameters that can be optimized to adjust the desired precision-recall balance, we selected a stringent set of peaks from the MACS2 peak calls that meet a −log_10_(FDR) threshold of greater than 10, and recalculated F1 scores in comparison with SEACR. Although the more stringent MACS2 peak calls had improved performance between 10M and 25M fragments, performance suffered at fragment subsampling levels below 10M reads, rendering SEACR superior at those levels (Figure 7—figure supplement 1B). Therefore, SEACR remains competitive with widely used ChIP-seq peak callers across multiple parameter selection strategies, even in the absence of arbitrary user input for the purposes of optimization. Although our conclusions are based on the presumption that high-scoring ENCODE peaks are true positives, the fact that they were called using MACS2 leads us to expect that the superior performance of SEACR on CUT&RUN data will generalize to any set of presumed true positives. Thus, SEACR is an optimal peak caller for CUT&RUN data across a range of read depths, and maintains a high percentage of true positive peak calls at low read depth.

Uniform peak calling is often confounded by the diverse distributions of chromatin proteins and modifications; for instance, transcription factors are expected to cluster at narrow genomic loci and adopt a typical “peaked” data structure, whereas many histone modifications such as H3K27me3 cover broad regions that are not easily summarized by the same methods that detect typical peaks. Since our signal block approach is agnostic to region width, we reasoned that SEACR might be equally successful at identifying broad domains as the peaks identified from Sox2, FoxA2, and H3K4me2 data. To test this, we called peaks using SEACR, MACS2 and HOMER (using “stringent”, “broad”, and “histone” settings, respectively) from an H3K27me3 CUT&RUN dataset (15) that contains broad domains by visual inspection. Remarkably, though SEACR called many fewer enriched regions than MACS2 or HOMER (28803, 97247, and 104524, respectively), SEACR regions covered more sequence (31.4 Mb) than either (28.1Mb and 18.3 Mb), indicating that SEACR regions are broader. Indeed, the average width of SEACR regions exceeded that of MACS2 and HOMER by nearly an order of magnitude (Figure 8A). Visual inspection of loci with broad H3K27me3 domains such as the HOXD cluster indicates that, whereas MACS2 and HOMER partition the domain into several subregions, SEACR maintains the majority of the domain structure in a limited number of large signal blocks (Figure 8B). These data indicate that SEACR is a promising tool for identifying large domains in CUT&RUN data in addition to spatially restricted binding sites.

**Figure 8.**
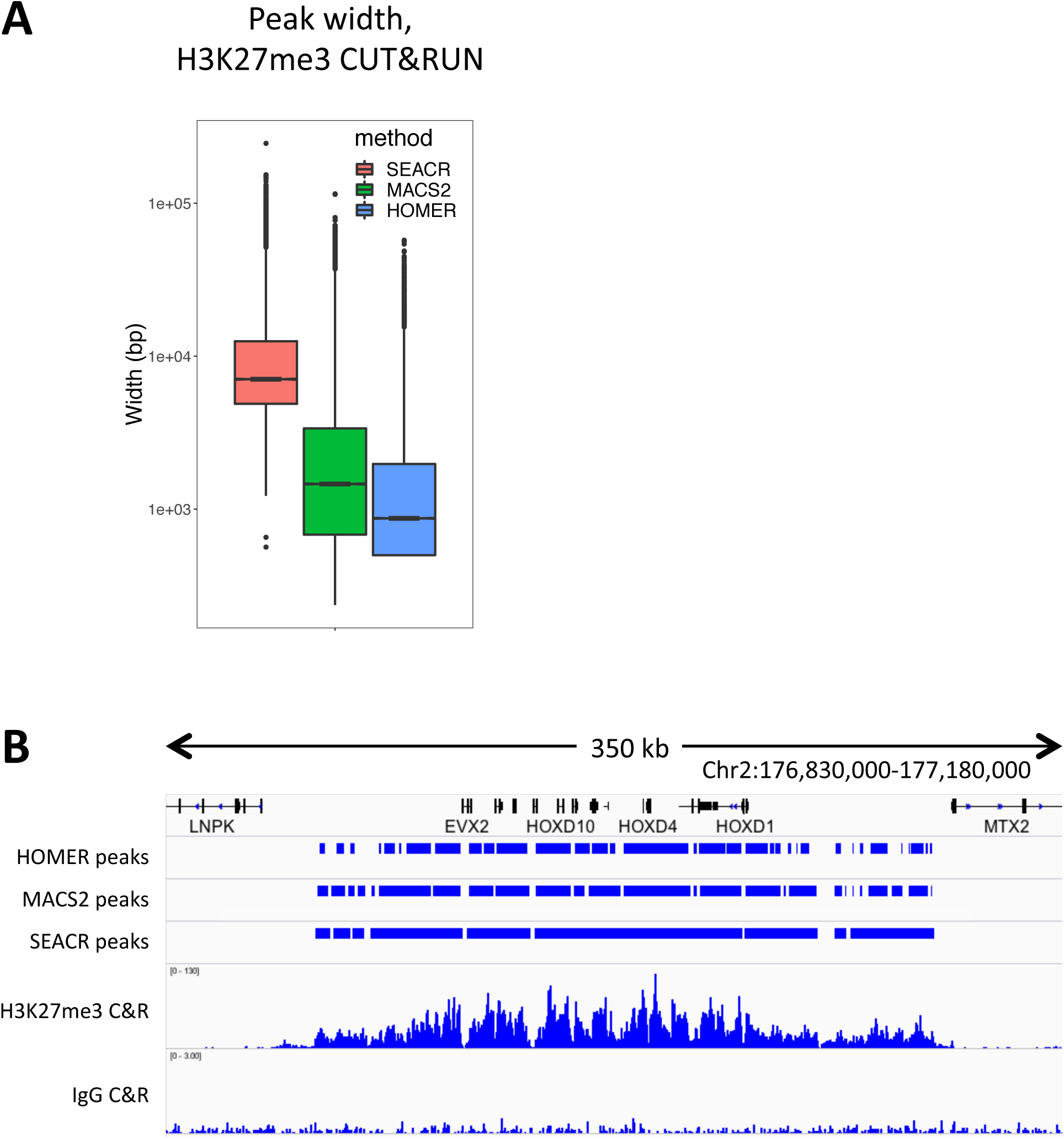
SEACR identifies coherent broad domains. **A**) CUT&RUN domains identified by SEACR are broader than those found by MACS2 or HOMER. **B**) A representative region of a H3K27me3 profile is shown. SEACR domains are relatively coherent compared with HOMER and MACS2 domains, which are fragmented into mixtures of wide and narrow segments.

## Conclusions

Since its introduction in our original *eLife* paper (1), the advantages of CUT&RUN over ChIP-seq has led to its rapid adoption, including publication of new CUT&RUN protocols for low cell numbers (14, 23), for plant tissues (25) and for high-throughput (15). The new CUT&RUN advances that we describe here are likely to be useful when applied in all of these protocols. Our improved CUT&RUN fusion construct simplifies reagent purification and eliminates the requirement for a secondary antibody against mouse primary antibodies. Our high-calcium/low-salt protocol minimizes time-dependent variability. Our discovery that carry-over *E. coli* DNA almost perfectly correlates with an added spike-in upgrades a contaminant to a resource that can be used as a spike-in calibration proxy, even *post-hoc* simply by counting reads mapping to the *E. coli* genome in existing CUT&RUN datasets.

We have also introduced a novel peak-calling strategy that takes advantage of the precise position and fragment spanning information that is present in CUT&RUN data. Popular peak-calling programs were designed around ChIP-seq data, where fragment spans are lacking owing to the widespread use of sonication and single-end sequencing. In contrast, our SEACR algorithm finds peaks in CUT&RUN data with a better precision/recall trade-off than the most popular ChIP-inspired peak-callers. The near-absence of false positives called by SEACR for Sox2 and FoxA2 transcription factors in cells that do not express them confirms the very high accuracy of CUT&RUN, in contrast to ChIP-seq, where reports of “Phantom Peaks” and other issues undermine confidence in peak calls (4-6). SEACR is also likely to be useful for other epigenomic datasets that capture fragment position and length information with high signal-to-noise. We expect that as the value of precise fragment information becomes better appreciated, for example in inferring chromatin dynamics (47), our block aggregation strategy will become increasingly powerful.

## Materials and methods

### Key Resources Table

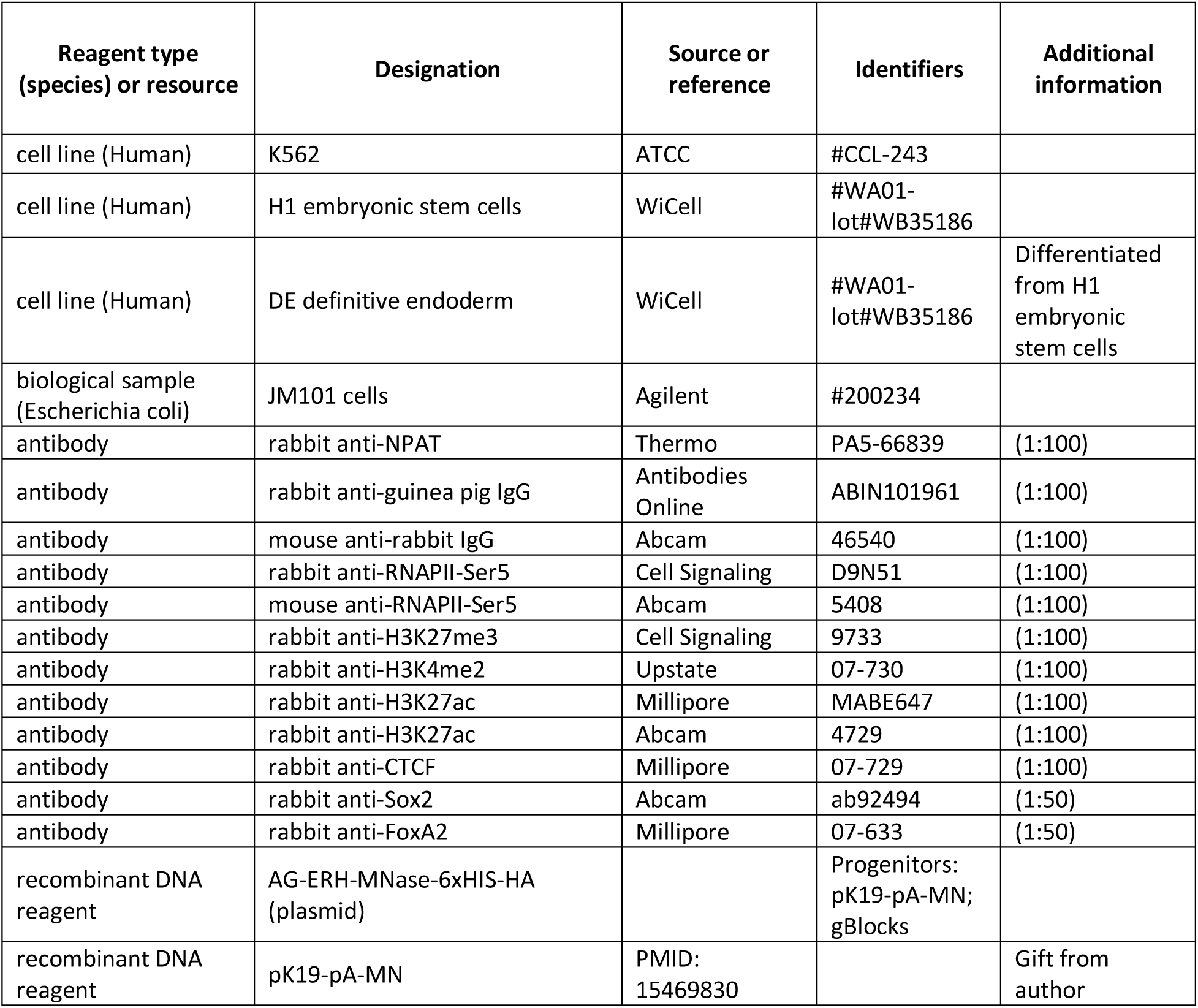
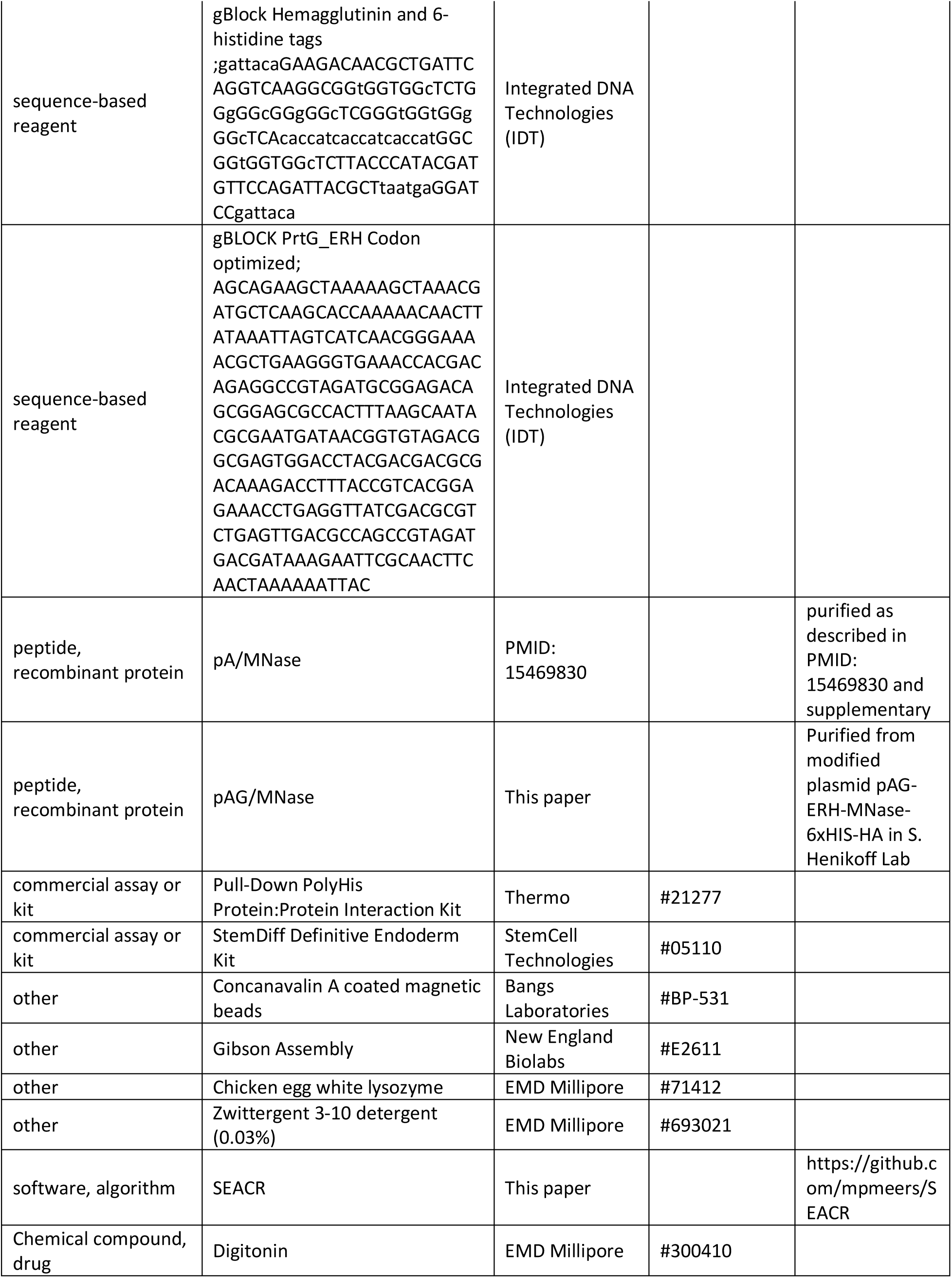
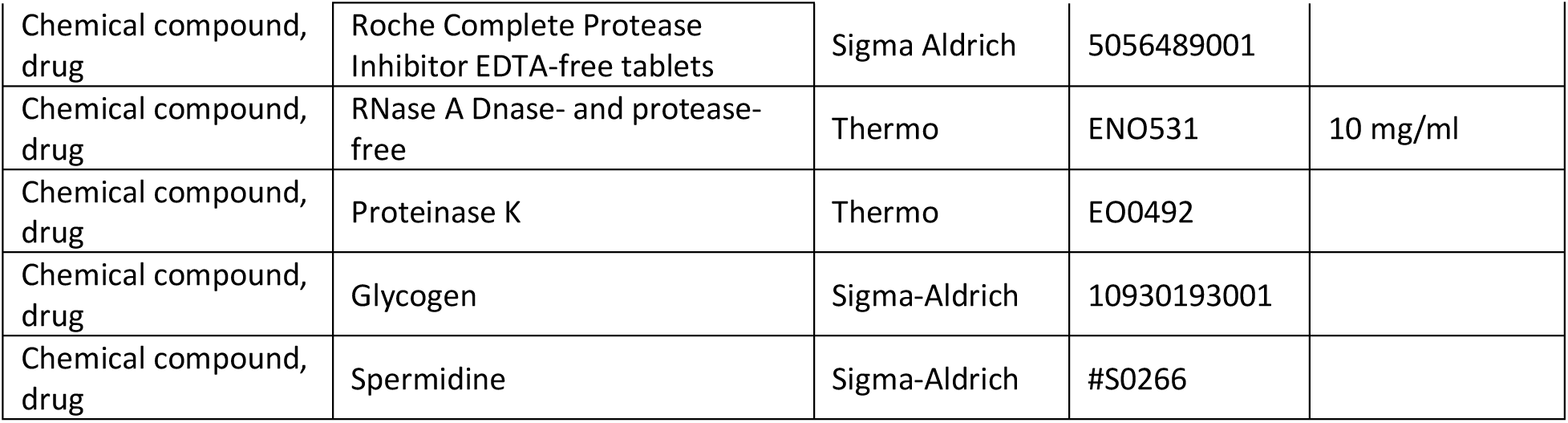

### Cell culture

K562 cells and H1 hESCs were cultured as previously described (15). Definitive endoderm (DE) was derived from a culture of H1 hESCs using the StemDiff Definitive Endoderm kit (StemCell Technologies).

### Construction and purification of an improved IgG-affinity/MNase fusion protein

Hemagglutinin and 6-histidine tags were added to the carboxyl-terminus of pA-MNase (13) using a commercially synthesized dsDNA fragment (gBlock®) from Integrated DNA Technologies (IDT), which contains the coding sequence for both tags, glycine-rich flexible linkers and includes restriction sites for cloning. Another IDT gBlock® containing the optimized protein-G coding sequence and homologous flanking regions to the site of insertion, was introduced via PCR overlap extension using Gibson Assembly® Master Mix (New England Biolabs cat. #E2611), following the manufacturer’s instructions. The sequence-verified construct was transformed into JM101 cells (Agilent Technologies cat. #200234) for expression, cultured in NZCYM-Kanamycin (50 µg/ml) and induced with 2 mM Isopropyl β-D-1-thiogalactopyranoside following standard protein expression and purification protocols. The cell pellet was resuspended in 10 ml Lysis Buffer, consisting of 10mM Tris-HCL pH 7.5, 300mM NaCl, 10mM Imidazole, 5mM beta-mercaptoethanol, and EDTA-free protease inhibitor tablets at the recommended concentration (Sigma-Aldrich cat. #5056489001). Lysis using chicken egg white lysozyme (10 mg/mL solution, EMD cat. #71412 solution) was followed by sonication with a Branson Sonifier blunt-end adapter at output level 4, 45 seconds intervals for 5-10 rounds or until turbidity was reduced. The lysate was cleared by high-speed centrifugation and purified over a nickel-agarose column, taking advantage of the poly-histidine tag for efficient purification via immobilized metal affinity chromatography. Cleared lysate was applied to a 20 ml disposable gravity-flow column 1.5 ml (0.75 ml bed volume) of NI-NTA agarose (Qiagen cat. #30210), washed twice in three bed volumes of Lysis Buffer. Lysate was applied followed by two washes at five bed volumes of 10 mM Tris-HCl pH 7.5, 300 mM NaCl, 20 mM Imidazole, 0.03% ZWITTERGENT 3-10 Detergent (EMD Millipore cat. #693021) and EDTA-free protease inhibitor tablets. Elution was performed with 1 ml 10 mM Tris-HCl pH 7.5, 300 mM NaCl, 250 mM Imidazole and EDTA-free protease inhibitor tablets. Eluate was dialyzed twice against a 750 ml volume of 10mM Tris-HCl pH 7.5, 150 mM NaCl, 1 mM EDTA, 1 mM PMSF to remove imidazole. Glycerol was then added to 50%, aliquots stored at –80 °C for long term storage and –20 °C for working stocks.

For purification, we used either the nickel-based protocol or the Pierce Cobalt kit (Pull-Down PolyHis Protein:Protein Interaction Kit cat. #21277 from Thermo Fisher). Similar results were obtained using either the nickel or cobalt protocol, although the cobalt kit alleviated the need for a sonicator, using a fifth of the starting material from either fresh culture or a cell pellet frozen in lysis buffer, and yielded more protein per volume of starting material. With the cobalt kit, 20 ml of culture yielded ∼100 µg of fusion protein.

### CUT&RUN using high-calcium/low-salt digestion conditions

Log-phase cultures of K562 cells were harvested, washed, and bound to activated Concanavalin A-coated magnetic beads, then permeabilized with Wash buffer (20 mM HEPES, pH7.5, 150 mM NaCl, 0.5 mM spermidine and a Roche complete tablet per 50 ml) containing 0.05% Digitonin (Dig-Wash) as described (14). The bead-cell slurry was incubated with antibody in a 50-100 µL volume for 2 hr at room temperature or at 4°C overnight on a nutator or rotator essentially as described (14). In some experiments, cells were permeabilized and antibody was added and incubated 2 hr to 3 days prior to addition of ConA beads with gentle vortexing; similar results were obtained (*e.g.* Figure 2B-D), although with lower yields. After 2-3 washes in 1 ml Dig-wash, beads were resuspended in 50-100 µL pA/MNase or pAG/MNase and incubated for 1 hr at room temperature. After 2 washes in Dig-wash, beads were resuspended in low-salt rinse buffer (20 mM HEPES, pH7.5, 0.5 mM spermidine, a Roche mini-complete tablet per 10 ml and 0.05% Digitonin). Tubes were chilled to 0°C, the liquid was removed on a magnet stand, and ice-cold calcium incubation buffer (3.5 mM HEPES pH 7.5, 10 mM CaCl_2_, 0.05% Digitonin) was added while gently vortexing. Tubes were replaced on ice during the incubation for times indicated in each experiment, and within 30 seconds of the end of the incubation period the tubes were replaced on the magnet, and upon clearing, the liquid was removed, followed by immediate addition of EGTA-STOP buffer (170 mM NaCl, 20 mM EGTA, 0.05% Digitonin, 20 µg/ml glycogen, 25 µg/ml RNase A, 2 pg/ml *S. cerevisiae* fragmented nucleosomal DNA). Beads were incubated at 37°C for 30 min, replaced on a magnet stand and the liquid was removed to a fresh tube and DNA was extracted as described (14). A detailed step-by-step protocol is available at https://www.protocols.io/view/cut-run-targeted-in-situ-genome-wide-profiling-witn6wdhfe/abstract. Extraction of pellet and total DNA was performed essentially as described (1, 16).

### DNA sequencing and data processing

The size distribution of libraries was determined by Agilent 4200 TapeStation analysis, and libraries were mixed to achieve equal representation as desired aiming for a final concentration as recommended by the manufacturer. Paired-end Illumina sequencing was performed on the barcoded libraries following the manufacturer’s instructions. Paired-end reads were aligned using Bowtie2 version 2.2.5 with options: --local --very-sensitive-local --no-unal --no-mixed --no-discordant --phred33 -I 10 -X 700. For MACS2 peak calling, parameters used were macs2 callpeak – t input_file –p 1e-5 –f BEDPE/BED(Paired End vs. Single End sequencing data) –keep-dup all –n out_name. Some datasets showed contamination by sequences of undetermined origin consisting of the sequence (TA)_n_. To avoid cross-mapping, we searched blastn for TATATATATATATATATATATATAT against hg19, collapsed the overlapping hits into 34,832 regions and intersected with sequencing datasets, keeping only the fragments that did not overlap any of these regions.

### Evaluating time-course data

If digestion and fragment release into the supernatant occur linearly with time of digestion until all fragments within a population are released, then we expect that CUT&RUN features will be linearly correlated within a time-course series. For CTCF, features were significant CTCF motifs intersecting with DNAseI hypersensitive sites (1). For H3K27Ac and H3K4me2, we called peaks using MACS2 and calculated the Pearson correlation coefficients between time points, displayed as a matrix of R^2^ values, using the following procedure:

1. Aligned fastq files to unmasked genomic sequence using Bowtie2 version 2.2.5 to UCSC hg19 with parameters: --end-to-end --very-sensitive --no-mixed --no-discordant -q --phred33 -I 10 -X 700.
2. Extracted properly paired read fragments from the alignments and pooled fragments from multiple samples.
3. Compared pooled fragments with (TA)_n_ regions of hg19 and kept those fragments that did NOT overlap any (TA)_n_ region using bedtools 2.21.0 with parameters: intersect -v -a fragments.bed -b TATA_regions.bed > fragments_not_TATA.bed.
4. Found peaks using macs2 2.1.1.20160309 with parameters: callpeak -t fragments_not_mask.bed -f BED -g hs --keep-dup all -p 1e-5 -n not_mask –SPMR.
5. Made scaled fractional count bedgraph files for each sample from bed files made in step 2. The value at each base pair is the fraction of counts times the size of hg19 so if the counts were uniformly distributed the value would be 1 at each bp.
6. Extracted bedgraph values for ±150 bps around peak summits for IgG sample and computed their means, which resulted in one mean score per peak.
7. Removed peaks from macs2 results in step 4 if the mean score was greater than the 99th percentile of all IgG scores to make a subset of the peaks lacking the most extreme outliers.
8. Extracted bedgraph values for ±150 bps around the subset of peak summits from step 7 for all samples and computed their means, which resulted in a matrix with columns corresponding to samples and one row per peak.
9. Computed correlations of matrix in 8 using R 3.2.2 cor(matrix, use=“complete.obs”) command.

### SEACR: Sparse Enrichment Analysis for CUT&RUN

SEACR was designed to call enriched regions from sparse CUT&RUN data, in which background is dominated by “zeroes” (i.e. regions with no read coverage). SEACR takes as input the following five fields: 1) Target data bedgraph file in UCSC bedgraph format (https://genome.ucsc.edu/goldenpath/help/bedgraph.html) that omits regions containing 0 signal; 2) Control (IgG) data bedgraph file; 3) “norm” denotes normalization of control to target data, “non” skips this behavior; 4) “relaxed” forces implementation of a maximum signal threshold in addition to the total signal threshold, and corresponds to the “relaxed” mode described in the text, whereas “stringent” avoids this behavior, and corresponds to “stringent” mode; 5) Prefix for output file.

Briefly, for each input bedgraph, we concatenated each region with adjacent regions to generate “signal blocks” that span all concatenated component regions, and calculated total signal for each signal block by taking the sum across all component regions of the region length (region end minus region start) multiplied by its bedgraph signal (column 4):

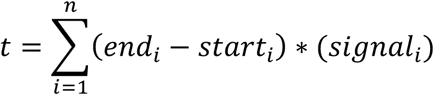

We designated maximum signal for each signal block as the maximum bedgraph signal value for any component region contained in the block. For normalization, we generated total signal density plots for all signal blocks from target and control data, identified the total signal AUC values that corresponded to the density peak of each plot, and multiplied total signal values for all control signal blocks by a “scaling factor” calculated by dividing the density peak total signal value for target data by the density peak total signal value for control data. To determine the total signal threshold *t*, we identified the value corresponding to the maximum value of *F* for the following function:

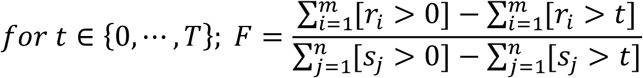

Where *T* is the maximum total signal value in any signal block, *r*_*i*_ is the total signal for an element *i* in the set of *m* target signal blocks, *s*_*j*_ is the total signal for an element *j* in the set of *n* target or control signal blocks, and *F* is the fraction of target signal blocks divided by total signal blocks remaining above threshold *t*. Once *t* is established, all target signal blocks exceeding *t* that do not overlap a control signal block that also exceeds *t* are retained. For “relaxed” mode, a similar threshold calculation is implemented for maximum signal, and any target signal blocks exceeding *t* for maximum signal that don’t overlap control blocks exceeding *t* for total signal are also retained.

To improve SEACR performance at high read depth as presented in Figure 7—figure supplement 1A, an additional bedgraph signal filtering step was added prior to determining signal thresholds. Briefly, for each integer *i* multiple of the smallest maximum signal recorded in the bedgraph *m*, we converted all bedgraph regions with signal less than or equal to *i*m* to 0 and quantified the percentage of the reference genome represented by 0 signal. We then repeated the 0 conversion for the minimum threshold *i*m* for which the percentage of the reference genome represented by 0 signal was at least 50%, and carried out SEACR.

MACS2 peaks were called using macs2 callpeak -f BEDPE --keep-dup all, with treatment and control files. For H3K27me3, the --broad flag was added. HOMER peaks were called by generating tag directories for target and control datasets, then using findPeaks, -style factor for TFs or H3K4me2, and -style histone for H3K27me3. For recall and precision analysis, we used ENCODE H3K4me2 ChIP-seq peaks (ENCFF099LMD) meeting a –log10(FDR) cutoff of 10 as the test set, and used bedtools intersect with the –u flag to calculate the percentage of test set regions overlapped (fraction of true positives) or the percentage of experimental peaks outside the test set (fraction of false positives). F1 scores were calculated as follows:

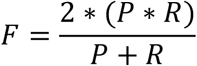

Where *P* is precision (fraction of called peaks that are true positives) and *R* is recall (fraction of true positives from test set identified).

## Availability

The plasmid pAG-ERH-MNase-6xHIS-HA is available from Addgene. Sequencing datasets are available from GEO (GSE126612). All scripts associated with SEACR are provided at https://github.com/mpmeers/SEACR.

## Acknowledgements

We thank Jorja Henikoff for data processing and analysis, and Christine Codomo and Tayler Hentges for technical support. We also thank all members of the Henikoff lab for valuable discussions and Kami Ahmad, Brian Freie and Bob Eisenman for comments on the manuscript. This work was supported by the Howard Hughes Medical Institute, and a grant from the National Institutes of Health (4DN TCPA A093) and the Chan-Zuckerberg Initiative.

## Author contributions

M.P.M and S.H. designed and performed all profiling experiments. T.D.B. designed and produced the pAG/MNase fusion construct. M.P.M. developed algorithms and M.P.M. and S.H analyzed the data. M.P.M, T.D.B and S.H. wrote the manuscript.

---

**Figure 1 – figure supplement 1.**
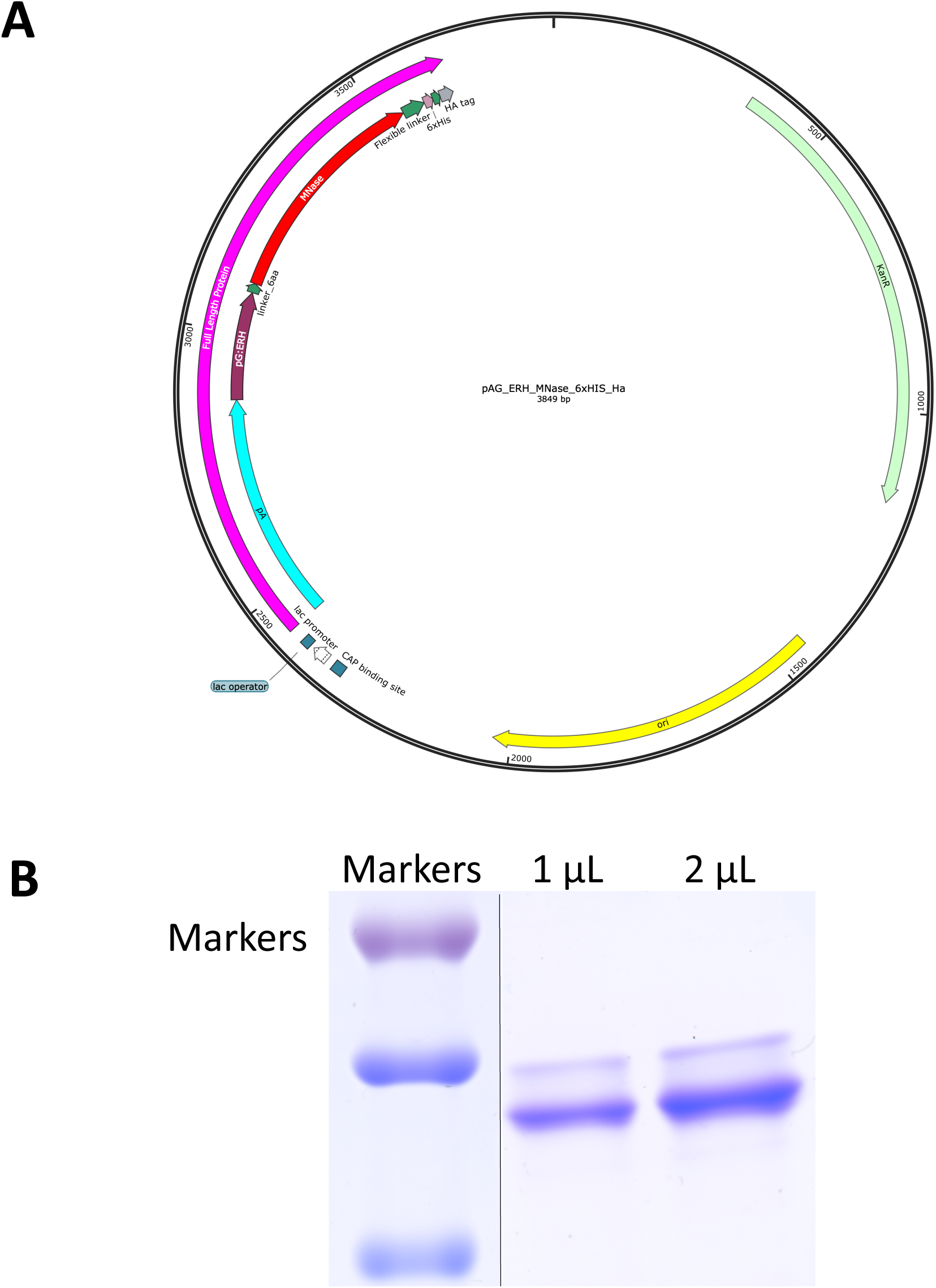
An improved fusion protein for CUT&RUN. **A**) Plasmid map of pAG-ERH-MNase-6xHIS-HA. **B**) Coomassie-stained gel of fusion protein eluted from nickel-agarose.

**Figure 1 – figure supplement 2.**
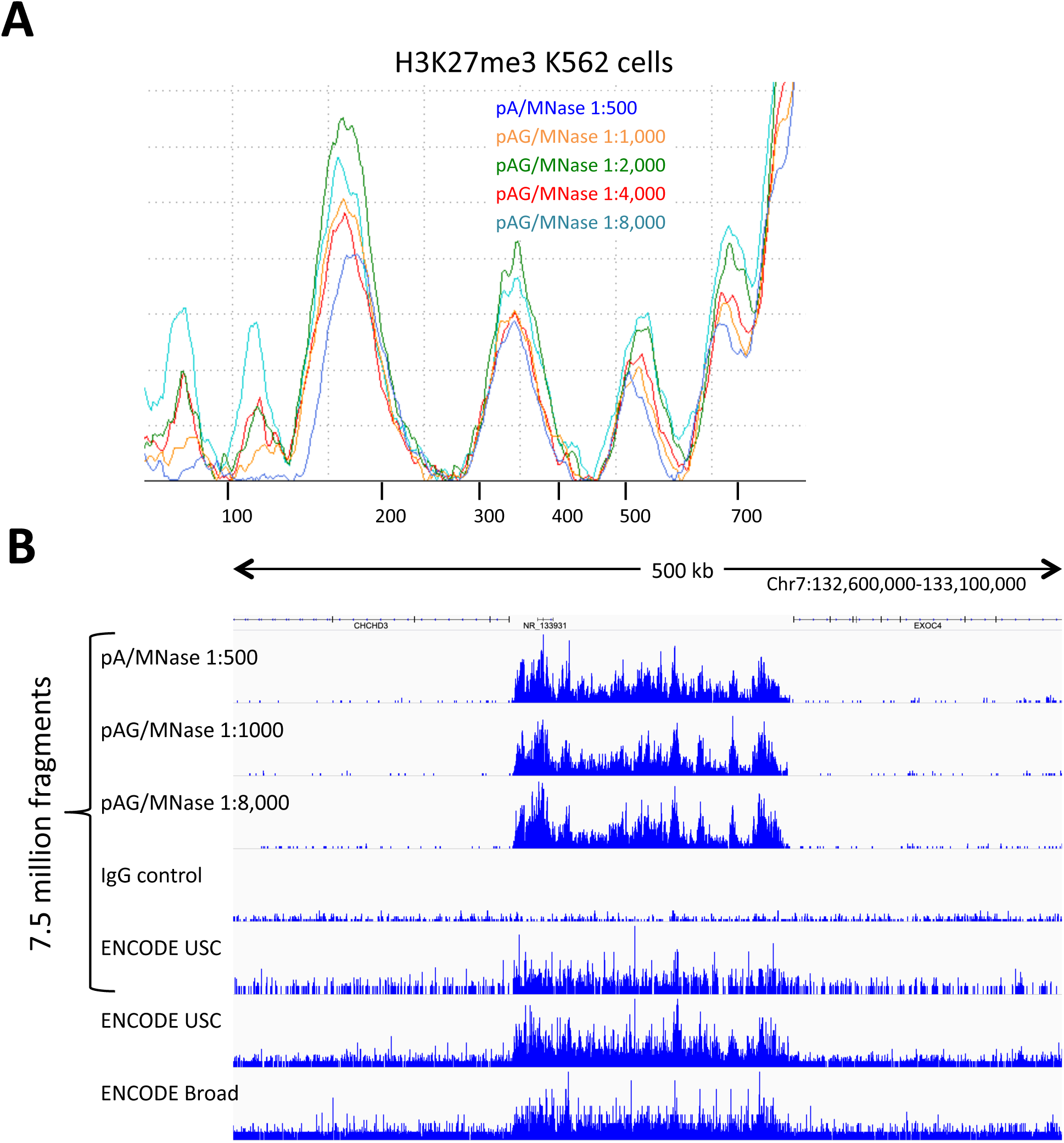
pAG/MNase titration. **A**) K562 cells were incubated with an antibody to H3K27me3 (CST #9733 Rabbit monoclonal), washed twice with 1 ml Dig-wash. The sample was split into aliquots for incubation with pA/MNase at the recommended concentration and a serial dilution of pAG/MNase, followed by 3 1 ml washes. After 30 min using the standard protocol, lImit digestions are seen at all dilutions for this abundant epitope, indicating that the amount of fusion protein used in this experiment was in excess. **B**) Representative tracks from these samples on the same normalized count scale show consistently low CUT&RUN backgrounds with excess pAG/MNase, which indicates that washes are sufficient to minimize non-specific background cleavages. ENCODE ChIP-seq tracks are shown for comparison, where USC used CST #9733, and Broad Institute used Millipore 07-449.

**Figure 4 – figure supplement 1.**
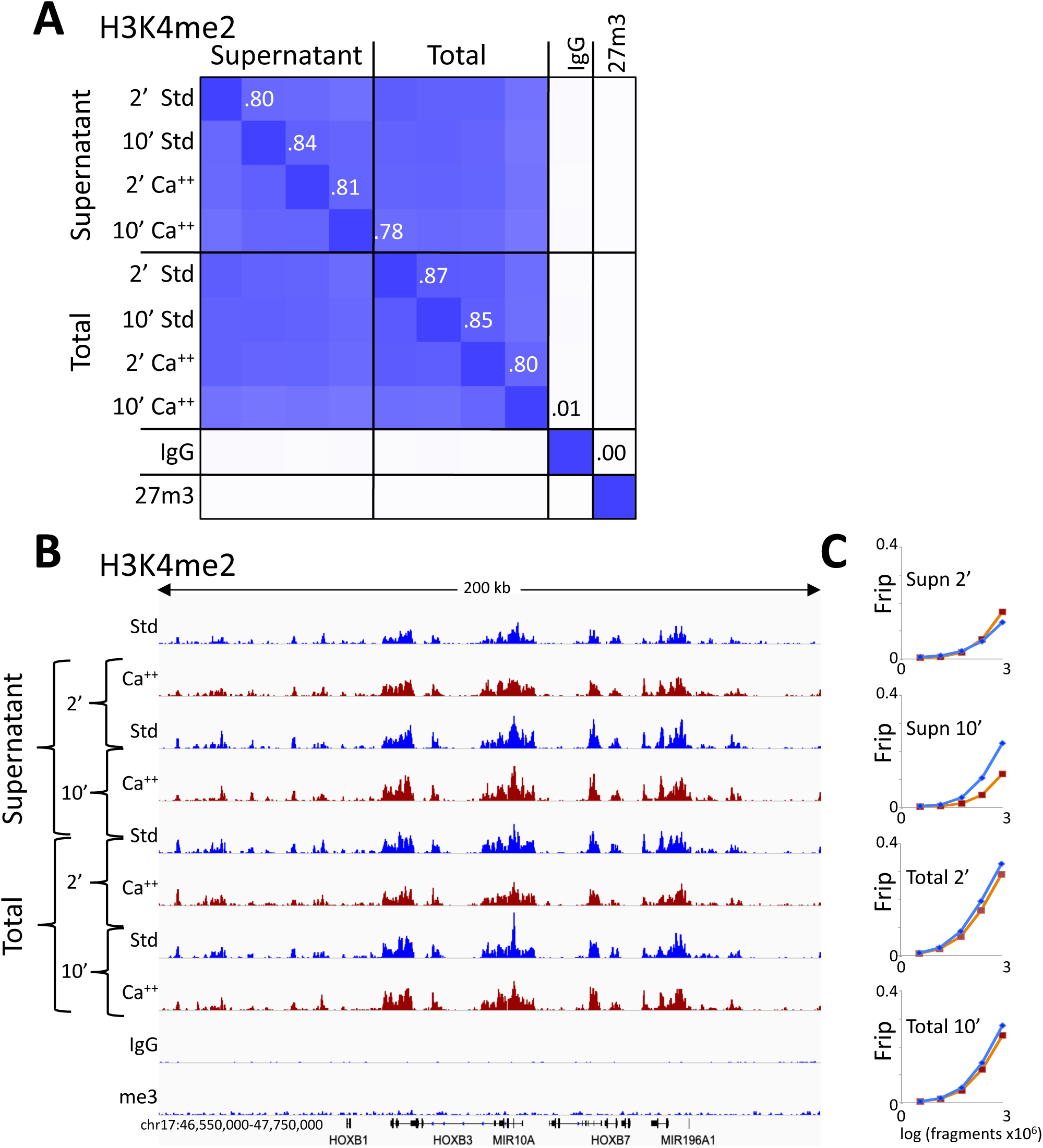
CUT&RUN consistency with high-Ca++/low salt digestion and total DNA extraction. **A**) H3K4me2 CUT&RUN time points with digestions using either the standard protocol or the high-calcium/low-salt protocol with either supernatant or total DNA extraction. To construct the correlation matrix, all 8 H3K4me2 datasets were pooled and MACS2 was used to call peaks, which yielded 64,156 peaks. Peak positions were scored for each dataset and correlations (R^2^ displayed with Java TreeView v.1.16r2, contrast = 1.25) were calculated between peak vectors. IgG and H3K27me3 (me3) negative controls were similarly scored. **B**) Same as Figure 4B. **C**) Same as Figure 4C.

**Figure 4 – figure supplement 2.**
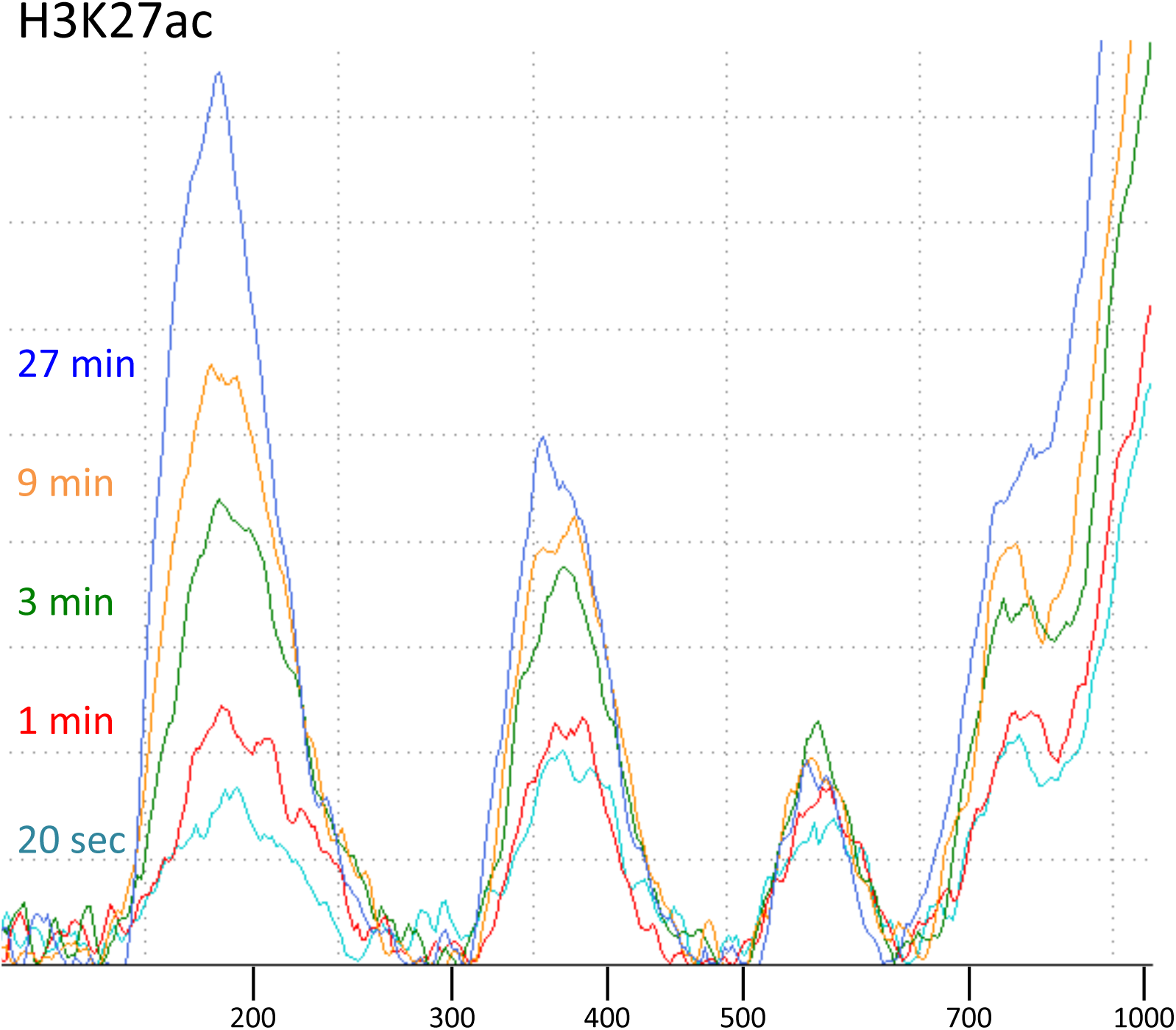
Tapestation analyses of an H3K27ac digestion time-course series. CUT&RUN with the low-salt/high-calcium protocol results in fragment release within 20 seconds at 0 °C.

**Figure 5 – table supplement 1.**
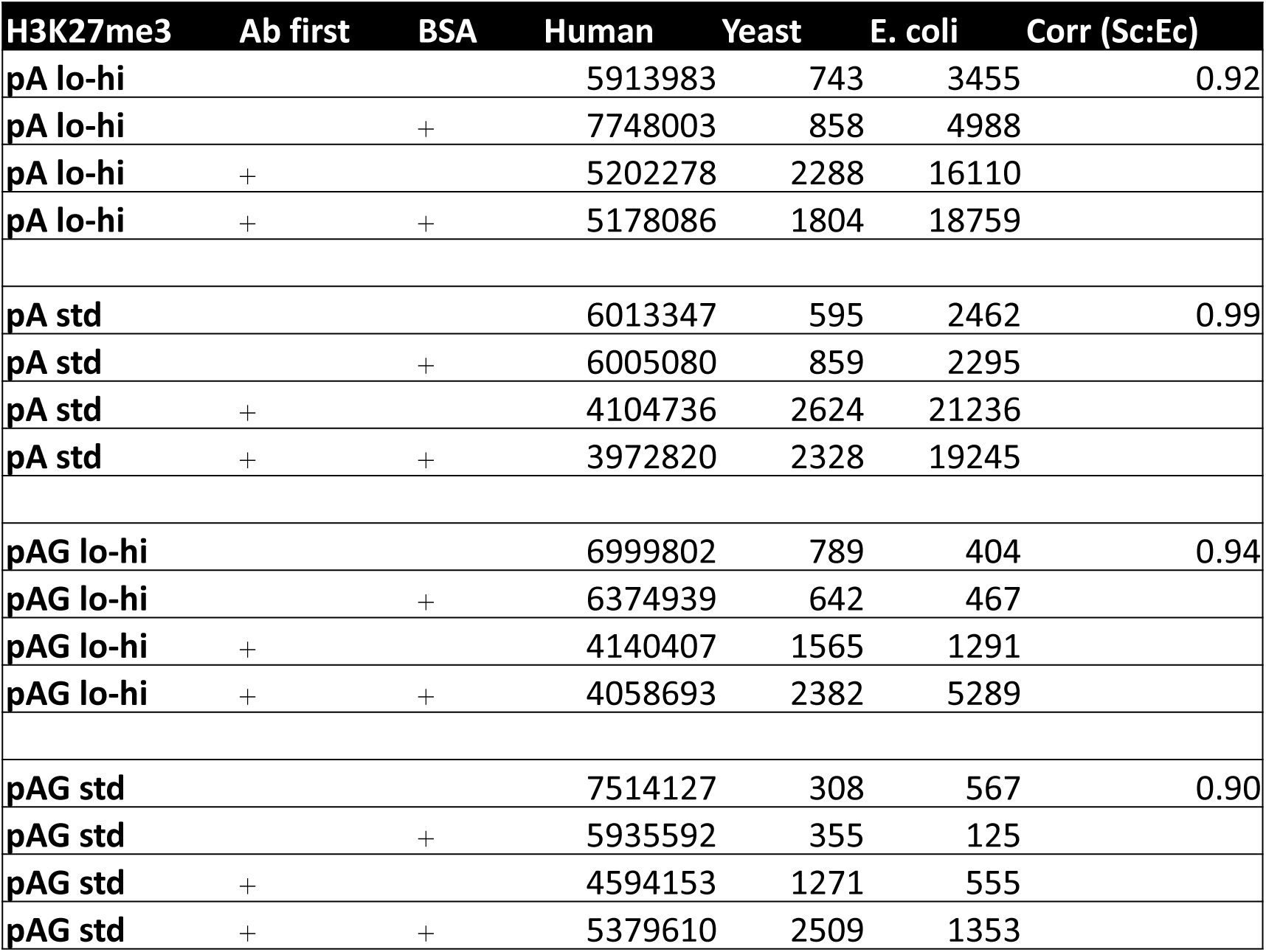
Carry-over *E. coli* DNA correlates closely with the heterologous spike-in for both fusion proteins and both low-salt/high-calcium and standard digestion conditions. CUT&RUN was performed for H3K27me3 in parallel for pA/MNase Batch #6 (pA), pAG/MNase (pAG) using both low-salt/high-calcium (lo-hi) and standard (std) CUT&RUN digestion conditions. Each sample started with ∼700,000 cells and 10 µL of bead slurry. Also varied in this experiment was addition of antibody followed by bead addition (Ab first) and addition of 0.1% BSA in the antibody buffer (BSA). Adding antibody first led to increased recovery of both yeast and E. coli DNA relative to human DNA, indicative of loss of cells prior to addition of fusion protein, possibly caused by loss of digitonin solubilization of membrane sugars.

**Figure 6 – figure supplement 1.**
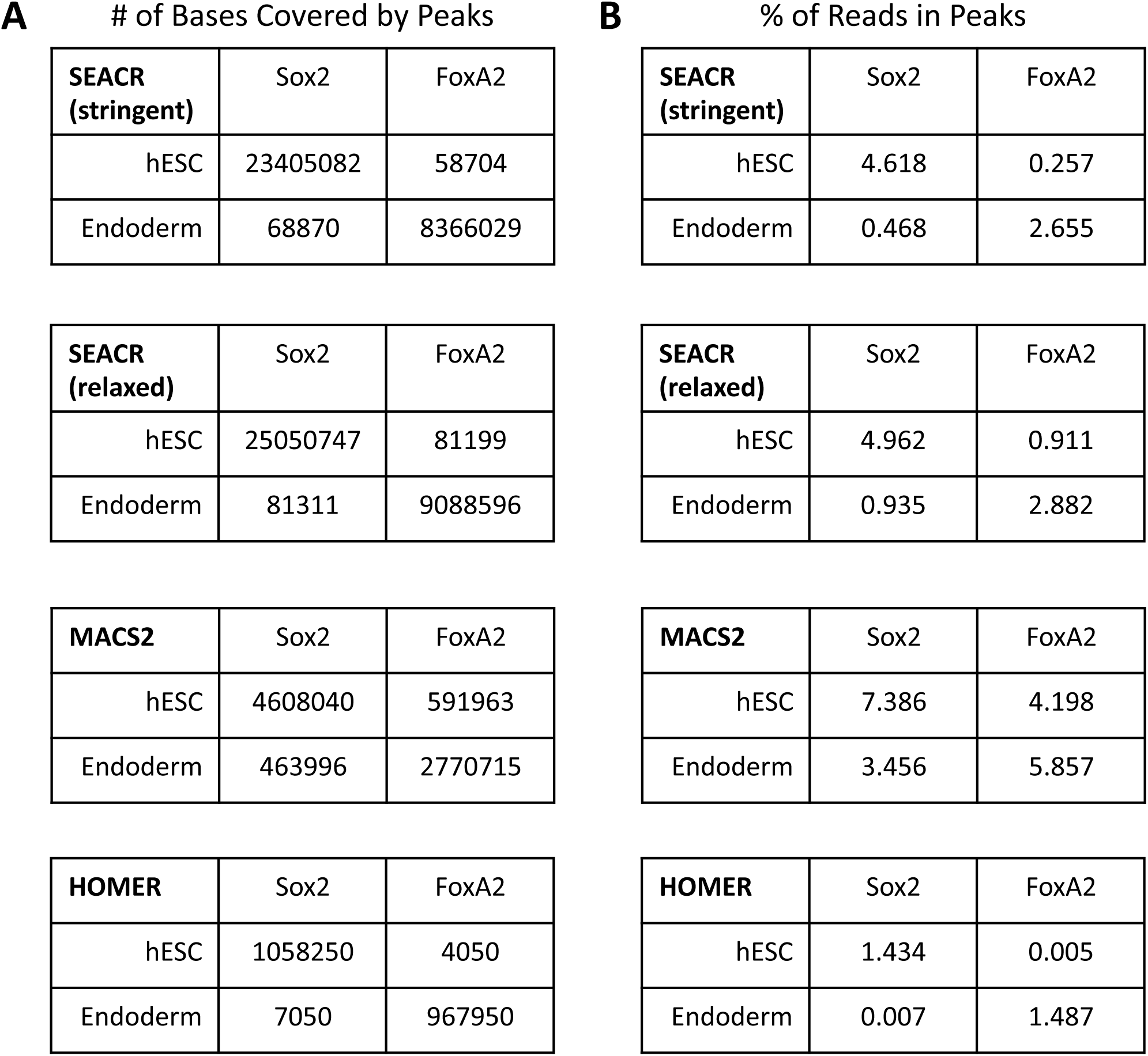
Peak calling metrics for SEACR. A-B) Table denoting the number of bases overlapped (A) or the percentage of reads in peaks (B) called by SEACR stringent mode (top row), SEACR relaxed mode (second row), MACS2 (third row), or HOMER (fourth row). Factor and cell type from which data are derived are indicated in columns and rows, respectively.

**Figure 7 – figure supplement 1.**
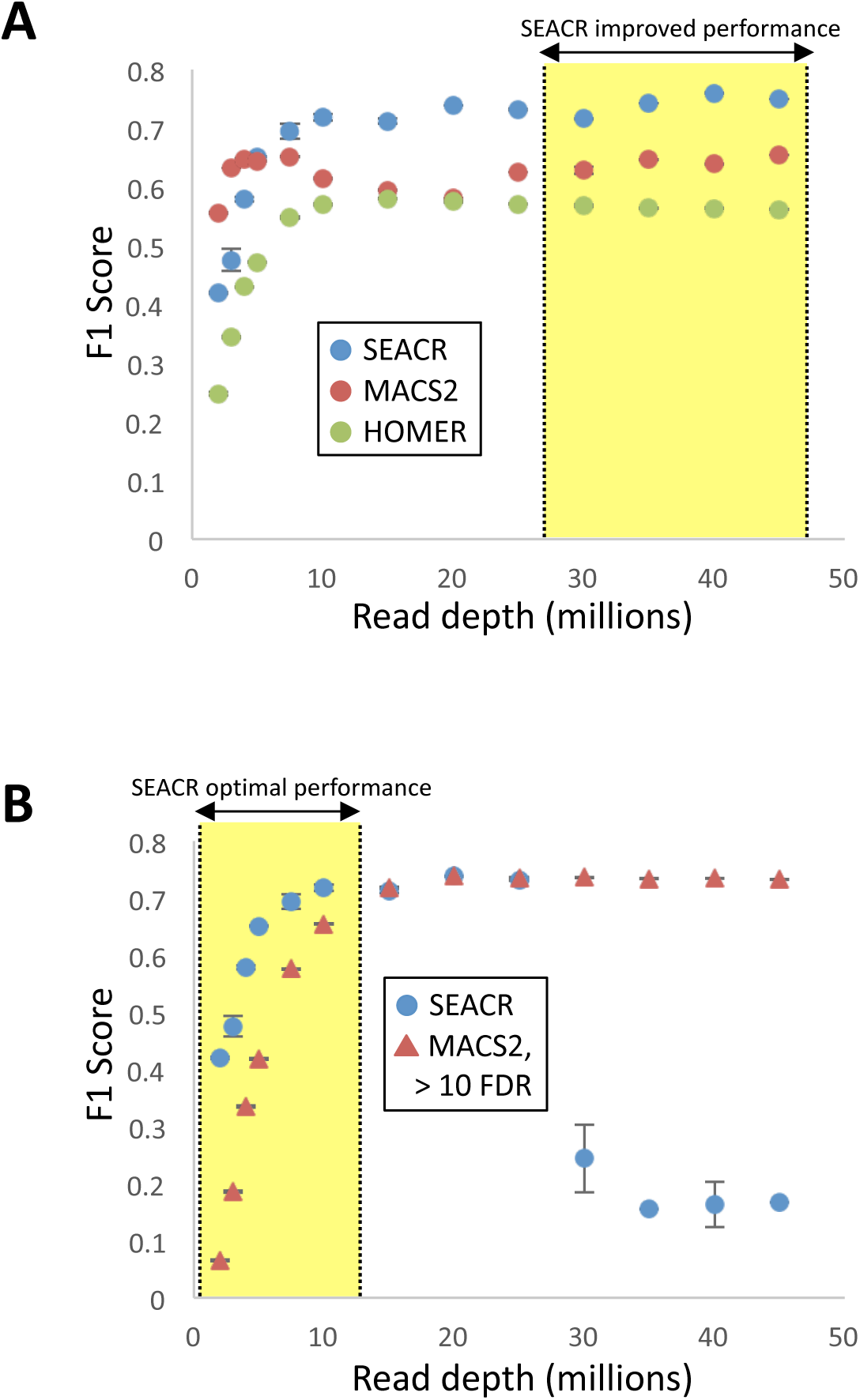
SEACR performance in comparison with other peak callers. A) F1 scores calculated as described for Figure 7D for SEACR using a genome coverage threshold, MACS2, and HOMER. Yellow shaded region indicates read depths at which SEACR performance improves with a genome coverage threshold relative to Figure 7D. B) F1 scores for SEACR and MACS2 peaks of greater than 10 –log10(FDR). Yellow shaded region indicates read depths at which SEACR performance is superior.

